# Long-range morphogen gradient formation by cell-to-cell signal propagation

**DOI:** 10.1101/2022.04.11.487794

**Authors:** Johanna E. M. Dickmann, Jochen C. Rink, Frank Jülicher

**Affiliations:** Max Planck Institute for the Physics of Complex Systems, Nöthnitzer Straße 38, 01187 Dresden, Germany; Max Planck Institute for Multidisciplinary Sciences, Am Faßberg 11, 37077 Göttingen, Germany; Cluster of Excellence, Physics of Life, TU Dresden, 01307 Dresden, Germany

**Keywords:** Morphogen gradients, tissue patterning, long-range patterning, cellular signaling, Wnt signaling, development, planarians

## Abstract

Morphogen gradients are a central concept in developmental biology. Their formation often involves the secretion of morphogens from a local source, that spread by diffusion in the cell field, where molecules eventually get degraded. This implies limits to both the time and length scales over which morphogen gradients can form which are set by diffusion coefficients and degradation rates. Towards the goal of identifying plausible mechanisms capable of extending the gradient range, we here use theory to explore properties of a cell-to-cell signaling relay. Inspired by the millimeter-scale Wnt-expression and signaling gradients in flatworms, we consider morphogen-mediated morphogen production in the cell field. We show that such a relay can generate stable morphogen and signaling gradients that emanate from a source of morphogen at a boundary. This gradient formation can be related to an effective diffusion and an effective degradation that result from morphogen production due to signaling relay. If the secretion of morphogen produced in response to the relay is polarized, it further gives rise to an effective drift. We find that signaling relay can generate long-ranged gradients in relevant times without relying on extreme choices of diffusion coefficients or degradation rates, thus exceeding the limits set by physiological diffusion coefficients and degradation rates. A signaling relay is hence an attractive principle to conceptualize long-ranged gradient formation by slowly diffusing morphogens that are relevant for patterning in adult contexts such as regeneration and tissue turn-over.

## 1. Introduction

Morphogens and morphogen gradients are key concepts in developmental biology. Morphogens are defined as secreted signaling molecules that spread through the tissue and specify cell fate in a concentration-dependent manner [1]. The decrease in morphogen concentrations as a function of distance from the site of secretion (the source) is referred to as morphogen gradient. The existence of morphogens has been predicted [2] as cited in [3], [4, 5] and studied theoretically, [4, 6, 7] decades before they were first observed experimentally [8, 9]. The current concept combines the ideas of ‘form producers’ introduced by Alan Turing [4], and the idea of positional information introduced by Lewis Wolpert [5] with a large body of experimental observations [8–19]. The emergence of the morphogen concept highlights the importance of theory in the elucidation and understanding of morphogenesis.

Morphogen gradients can form when morphogens are secreted in a local source and spread by diffusion through a target tissue where they are also degraded [6, 13, 15]. This model can account for observations of morphogen gradients in different model organisms including the fruit fly embryo [13–15, 20] and the zebrafish embryo [16, 17]. In these systems, morphogen gradients typically reach length scales in the range of tens to a few hundreds of micrometers [8, 13–17, 21, 22]. Salient biophysical parameters, i.e. diffusion coefficient and morphogen degradation rate, have been estimated in quantitative experiments [13–17, 22]. These measured parameter values can explain the formation of morphogen gradients spanning the observed tissue dimensions within the relevant developmental time interval. Tissue patterning by morphogen gradients does not only occur during development on the micrometer dimensions of early embryos, but can also occur at larger spatial dimensions of adult tissues. Examples for patterning of adult tissues by soluble signaling molecules include the organization of mammalian liver lobules [23, 24], axolotl regeneration [25], as well as regeneration and steady-state turn-over of planarian flatworms [26–32]. These examples show that signaling gradients are relevant to pattern tissues on millimeter length scales and possibly even up to centimeter length scales and thus require mechanisms capable of gradient formation over post-embryonic length scales.

In principle, morphogen gradients formed by diffusion and degradation can achieve arbitrary decay lengths, either by increasing the diffusion coefficient or by increasing the morphogen life time. However, constraints on parameter values (e.g., the diffusion coefficient of proteins in aqueous solution [22]) or the typical time scales of biological processes (e.g., observed growth or regeneration times) impose limits on the system’s dimensions in which gradient formation by diffusion and degradation is feasible. For instance, it has been suggested that extracellular diffusion of Fgf8 (*D* = 53±8*µ*m^2^*/*s [16]) together with the estimated degradation rate (*k* = *D/λ*^2^ = (1.3 ± 0.3) × 10^*−*3^ 1*/*s [16]) gives rise to an Fgf8 gradient with a decay length of *λ* = 197 ± 7 *µ*m [16] during embryonic development of the zebrafish on a time scale of around 13 min, thus matching the dimension of the embryonic structure to be patterned and the time scale relevant for development. However, many morphogens are lipid-modified [33–35] and therefore poorly soluble in the aqueous tissue environment, which results in slow diffusion [36–38]. This is in particular the case for Wnt [35] and it has been debated how far Wnt can spread by diffusion [36, 37, 39–43]. Already when assuming a diffusion coefficient one order of magnitude smaller than the above cited value for Fgf8, molecular life times of the order of months would be required for the generation of the millimeter-scale Wnt signaling gradient that organizes the main body axis in planarians [26–28, 31, 32]. This poses the fundamental question of what other means of gradient formation could be better suited to generate such long-raged gradients.

Directed molecular transport is one concept to increase the decay length of gradients that has been examined [44]. A directed motion of molecules through the system (drift) can result in faster and more long-ranged gradient formation as compared to diffusive spreading, see Appendix A. Drift could be caused by active intracellular transport processes, or by extracellular fluid flows, generated for example by coordinated cilia beating or by coordinated contractions [44]. An alternative mechanism for gradient formation has been introduced in the “bucket brigade” model by Kerszberg and Wolpert in 1998 [45] in which morphogens move between transmembrane receptors along and between cells. Receptors recently bound by signaling molecules become refractory, preventing “backward” spreading of the morphogen. Further, the morphogen can be handed over from receptors on one cell to those on another [45]. This leads to the formation of gradients of receptor-bound morphogens. Kerszberg and Wolpert briefly discuss whether the hand-over of morphogens between cells could be replaced by the idea of positive feedback in which new morphogens are generated in response to a morphogen binding to a receptor [45]. They conclude that this would prevent the morphogen concentration from decaying away from the local source and would thus not be a feasible concept of gradient formation [45].

In this paper, we revisit the idea of positive feedback as a means of gradient formation. Inspired by the evidence for Wnt-mediated Wnt expression and the ensuing Wnt expression gradients in planarians [30, 32, 46] we explore how tissue-scale morphogen gradients can form by signaling relays. We develop a model in which the extracellular morphogen gradient and resulting intracellular gradient of signaling activity are connected by a positive feedback loop (i.e. Wnt-mediated Wnt expression). This describes morphogen-mediated morphogen production and introduces a signaling relay by which morphogens can propagate in the tissue. In fact, the whole tissue effectively becomes a signaling-dependent source due to the positive feedback. A signaling-independent source of morphogen at the boundary initiates the signaling relay (inspired by the putative role of a tail-tip expressed Wnt ligand in planarians [32, 46]). In contrast to the work by Kerszberg and Wolpert [45], our results show that a signaling relay can induce stable morphogen and signaling gradients. Moreover, we show that spreading of the signal by a cell-to-cell relay leads to effective diffusion of the morphogen through the system. That is, individual molecules do not have to travel long distances but are instead produced anew further down the system in a feedback-dependent manner. Furthermore, it leads to an effective degradation rate smaller than the molecular degradation rate. This way, a cell-to-cell signaling relay can efficiently generate long-ranged signaling gradients for physiologically relevant molecular diffusion coefficients and degradation rates on relevant time scales. Furthermore, when this feedback-dependent production is combined with secretion polarity in the plane of the cell field, the cell-to-cell relay additionally leads to the emergence of an effective drift resembling directed transport of morphogens through the system in the absence of molecular transport. This can further increase the decay length of the signaling and morphogen gradients. Finally, the range of the gradients can be set by regulating the feedback strength, allowing the system to adjust gradient range without the need to change diffusivity or molecular life time of the morphogen. Altogether, this makes cell-to-cell relay an attractive concept to explore in systems with millimeter-scale patterns as observed during regeneration such as in planarians, hydra, zebrafish, and salamanders.

## 2. Gradient formation by cell-to-cell signaling relay

In this paper we focus on morphogen gradient formation by a cell-to-cell relay where individual cells transmit the signaling information to their neighbors. In particular, our approach takes the coupling between intracellular levels of signaling activity and the extracellular morphogen levels into account. This differs from scenarios where intracellular signaling gradients are strictly downstream of extracellular morphogen levels. To capture the coupling, we account for individual cells that react to the morphogen concentration in the extracellular space by increasing their intracellular signaling activity, captured as the concentration of an intracellular signaling molecule downstream of the morphogen. This increased signaling activity eventually leads to the production and secretion of more morphogen, increasing the extracellular morphogen concentration. This way, the information is relayed from cell to cell as the whole cell field becomes a signaling-dependent source. The level of intracellular signaling activity and concentration of extracellular morphogen are thus interdependent. The gradient is positioned and oriented by a localized, signaling-independent source of morphogen at the proximal boundary. From this signaling-independent source, the morphogen profile propagates via the cell-to-cell signaling relay towards the distal end.

The schematic of the cell-based model is shown in figure 1. We denote by *b*_*n*_ the signaling activity in cell *n*, where *n* = 0, …, *N* −1, and *N* is the number of cells along the length of the system. Thus, *b*_*n*_ denotes the concentration of an intracellular signaling molecule that is downstream of the morphogen. Similarly, we introduce the morphogen concentration in the extracellular space *a*_*n*_, where *n* refers to the extracellular space between the cells *n* − 1 and *n*. For simplicity we use a discrete description of the extracellular space with concentrations *a*_*n*_ averaged within a cell length. The dynamic equations for these concentrations read

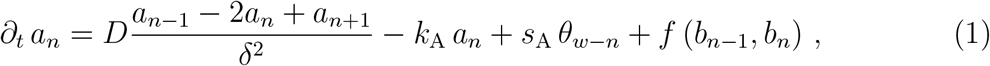

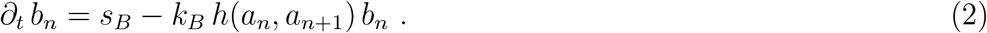

**Figure 1.**
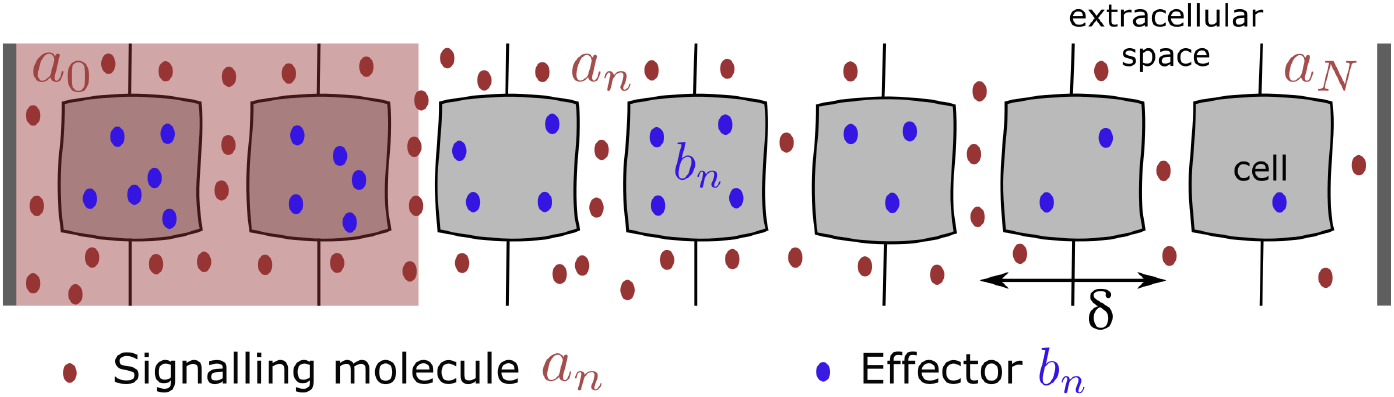
Schematic representation of the model geometry in one dimension. The extracellular morphogen *a*_*n*_ (depicted as red dots) is constantly produced in the signaling-independent source region shaded in red at the proximal end (*n* = 0). Moreover, it is produced throughout the system, which becomes a signaling-dependent source due to the signaling relay. It spreads across the extracellular spaces, where it is also degraded, towards the distal end (*n* = *N*). The intracellular signaling activity *b*_*n*_ (depicted as blue dots) is confined to the cell interior and the intracellular signaling molecules do not move between cells of size *δ*. The dynamics of the system is given by equations (1) and (2).

Here, *D* denotes the diffusion coefficient, *δ* the width of a cell, and *k*_A_ the rate of morphogen loss which could be due to degradation, internalization or leakage. For simplicity, we refer to *k*_A_ as the degradation rate of the morphogen. The rate of production in the signaling-independent source region (shaded in red in figure 1), is denoted *s*_A_. This signaling-independent source region at the proximal end (*n* = 0) of the system is described by the function *θ*_*w−n*_, where *θ*_*w−n*_ = 1 inside the source for *n < w*, and *θ*_*w−n*_ = 0 outside the source for *n > w*. At the source boundary, *n* = *w*, we use *θ*_*w−n*_ = *p* where *p* describes a cellular asymmetry of molecular secretion, see below. The distal end is at *n* = *N*. In equation (1), the positive feedback gives rise to an additional source *f* (*b*_*n−*1_, *b*_*n*_) that depends on the signaling activities in the two adjacent cells, *b*_*n−*1_ and *b*_*n*_. We refer to this as the signaling-dependent source. Equation (2) describes the dynamics of the signaling activity. Here, *s*_B_ denotes the rate of production of the intracellular signaling molecule, *k*_B_ is a degradation rate and *h*(*a*_*n*_, *a*_*n*+1_) describes the positive feedback regulating the degradation of the intracellular signaling molecule by the morphogen concentrations in the adjacent extracellular spaces *a*_*n*_ and *a*_*n*+1_. The boundary conditions are discussed in Appendix B. The signaling-dependent source *f*, as well as the regulation of the degradation of the intracellular signaling molecule *h* are given by

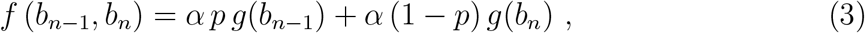

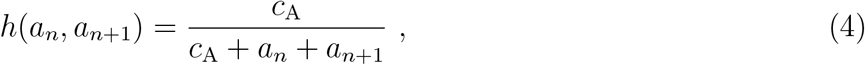

where *g*(*b*_*n*_) = *b*_*n*_*/*(*c*_B_ +*b*_*n*_) denotes the Hill activator function, and *c*_B_ is a Hill activation threshold, i.e. the concentration of *b*_*n*_ at which half-maximal activation is reached. Similarly, *h* is the Hill inhibitor function and *c*_A_ is the Hill inhibitor coefficient at which half-maximal inhibition is reached. Here, we use Hill exponents of 1 because we consider cells that respond linearly to weak stimuli before the response saturates at high stimuli. The maximal signaling-induced secretion rate is denoted *α*. Equation (3) describes polarized morphogen secretion. Such polarized secretion might for instance arise from distinct activities of the two lateral sides of the cell to secrete morphogen. This secretion polarity *p* is associated with cell polarity in the tissue akin to planar cell polarity. In the case of non-polar cells 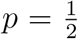. Extreme secretion asymmetries correspond to *p* = 0 and *p* = 1, respectively.

Equations (1) - (4) describe a positive feedback between morphogen concentrations and concentrations of the intracellular signaling molecule, where the parameter *α* sets the overall feedback strength. This feedback consists of two elements: Equation (3) contains a positive feedback element stimulating morphogen secretion with increasing intracellular signaling activity, the signaling-dependent source. Equation (4) also is a positive feedback element, increasing signaling activity for increased morphogen levels. This feedback element is positive due to inhibition of inhibition, a principle commonly observed in signaling pathways.

Our model is inspired by Wnt signaling. It is known that high extracellular Wnt concentrations prevent intracellular *β*-Catenin degradation [47] as captured by (2), corresponding to inhibition of inhibition. Equation (1) is motivated by observations in planarian flatworms suggesting that high intracellular *β*-Catenin levels lead to increased expression of several Wnt-signaling pathway components, including several Wnt ligands [32].

The cell-to-cell relay involving positive feedback is conceptually different from gradient formation by diffusion and degradation. The relay turns the whole system into a signaling-dependent source. Thus, individual signaling molecules do not have to move a distance spanning the entire gradient. They can form a long-ranged gradient even if they only move between neighboring cells. In the following, we analyze the impact of the positive feedback on the shape of the gradient and the dynamics of gradient formation.

## 3. Regulation of the gradient range by signaling feedback

### 3.1. Morphogen and signaling gradients at steady state

We first consider steady-state concentration profiles in order to discuss the range of the morphogen and signaling activity-gradients. We quantify the range of the gradients as the decay length *λ* over which they decay. Note that here we measure all lengths relative to the cell size *δ* and *λ* is therefore the number of cells corresponding to the gradient range. We can then analyze the impact of the positive feedback on the gradient range. The steady-state solutions to equations (1) and (2) are denoted 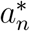 and 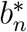 and obey 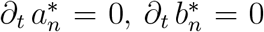. We can obtain these solutions numerically, see figure 2 and Appendix C. However, to discuss the impact of the feedback, we use an approximation which we can obtain analytically. Note that at steady-state, equation (2) implies that the profile of the intracellular signaling activity is linearly dependent on the morphogen concentration profile:

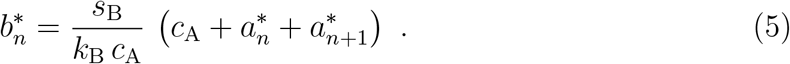

**Figure 2.**
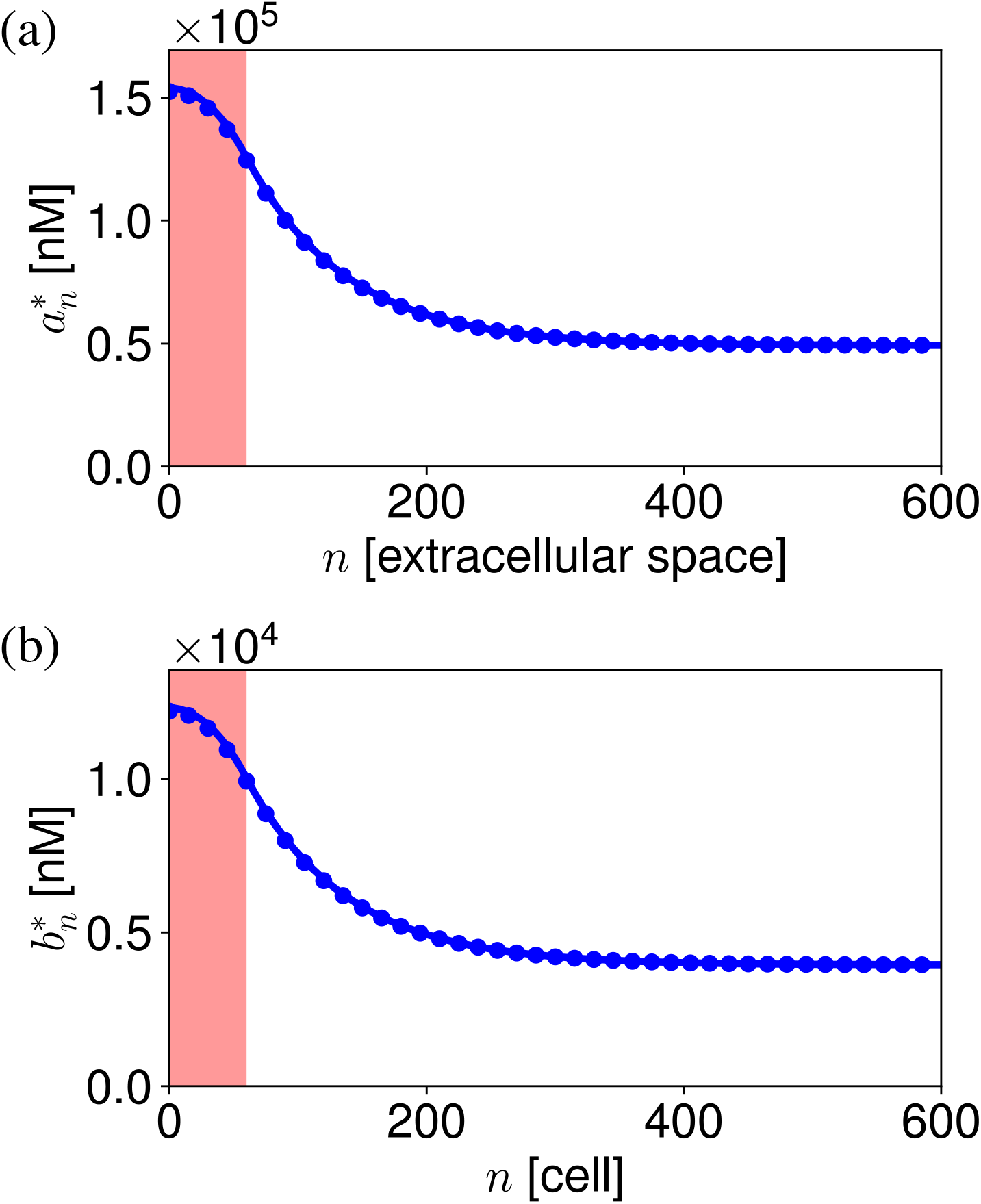
Steady-state concentration profiles of the morphogen (a) and the intracellular signaling activity (b). The signaling-independent source region is shaded in red. Circles denote the numerically obtained steady-state solution to the non-linear equations, lines indicate the analytical approximation to the steady state.

Thus, at steady state, the signaling activity-gradient can be obtained from the morphogen gradient using equation (5).

We focus our discussion on the morphogen gradient 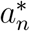 in the following. We obtain an approximation for the steady state by matching simplified solutions inside the source region with *n* = 0, … *w*, and outside of the source region with *n* = *w* + 1, …, *n*

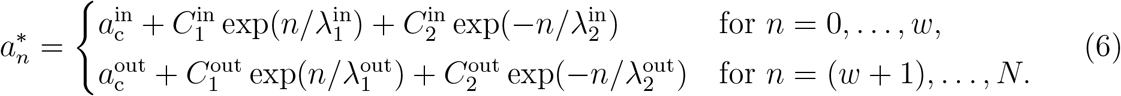

Here, the gradient amplitudes 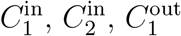, and 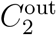 are determined by the boundary conditions, as well as the matching condition at the source boundary, see Appendix D. The decay lengths 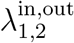 can be expressed explicitly, see Appendix D. We focus on the decay of the gradient outside the source, which occurs over a range 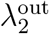. This term dominates for large system sizes outside of the source and therefore sets the gradient range

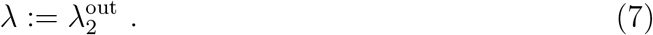

The decay length *λ* can be related to effective transport coefficients which we discuss in the next section.

### 3.2. Effective coefficients of molecular transport

The model for gradient formation by cell-to-cell signaling relay introduced in the last sections describes the kinetics of molecular transport and signaling at the scale of cells. In this model, a gradient with a long range emerges revealing that the system generates concentration patterns at larger scales. In the spirit of a hydrodynamic theory, molecular concentration profiles can be captured by an effective continuum description at scales much larger than cells. In this limit, it has the form of a convection-diffusion-degradation equation:

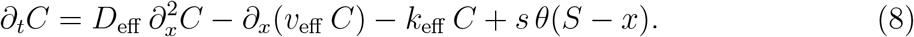

Here, *D*_eff_ is an effective diffusion coefficient, *k*_eff_ is an effective degradation rate, and the velocity *v*_eff_ describes an effective drift. Production with rate *s* is limited to a local source of width *S*, which corresponds to the signaling-independent source in the cell-based model. The spatial coordinate in the direction along which the gradient forms is denoted *x*, and *θ* denotes the Heavyside function, where *θ*(*y*) = 0 for *y <* 0 and *θ*(*y*) = 1 for *y* ≥ 0. In equation (8), 0 *< x* ≤ *L* with a tissue of size *L* and the source at the proximal boundary. Positive values of *v* indicate drift in positive *x*-direction. We can estimate effective transport coefficients as well as the effective degradation rate by comparing the large scale dynamics of our cell-based model to the corresponding behavior of the continuum model given by equation (8), see Appendix E. We obtain

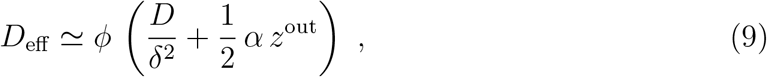

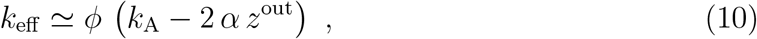

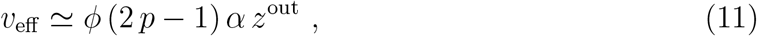

where

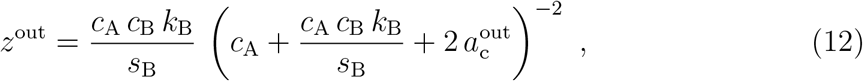

where 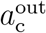 is given in Appendix D, see equation (D.3), and the coefficient *ϕ* is given in Appendix E, see equation (E.22). Note that, while the effective diffusion coefficient and degradation rate contain the respective molecular quantities modulated by the feedback strength *α*, the effective drift is a purely effective drift due to polarized secretion of molecules with polarity *p* in the complete absence of a molecular drift. That is, molecules appear to be transported through the system, when actually they are produced and secreted with secretion polarity *p*. Accordingly, the effective drift vanishes in the absence of secretion polarity at 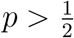.

Using these effective quantities, the decay length of the morphogen gradient at steady-state is given as:

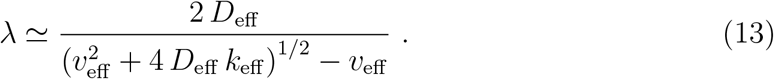

Based on this expression for the decay length, we can discuss the impact of the signaling relay on the gradient range, based on the effective quantities we identified above. We begin this discussion by focusing on the influence of the feedback strength *α*.

### 3.3. Regulation of gradient range by signaling-feedback strength

The positive feedback strength *α* can regulate the gradient range by modulating all effective quantities introduced above. In the absence of positive feedback, *α* = 0, the model becomes a diffusion/degradation system and thus the decay length simplifies to the well-known expression *λ* = *δ*^*−*1^(*D/k*)^1*/*2^ [15], see Appendix A, equation (A.5). In the presence of positive feedback, *α >* 0, the range of the gradient is increased as compared to the case *α* = 0, see figure 3. In particular, the decay length exhibits a maximum near a feedback strength *α*^crit^. Here, *α*^crit^ corresponds to a critical point that occurs for a linear feedback response when *g*(*b*_*n*_) is replaced by a linear function of *b*_*n*_. In this case, the concentration grows without bounds for *α > α*^crit^ with

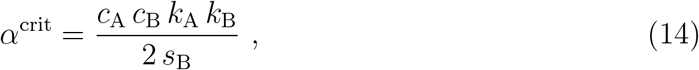

and no steady state exists. Such a linear regime occurs in the limit when the Hill activation threshold *c*_B_ is much larger than the signaling activity *b*_*n*_. In this case, *g* becomes *g*_lin_ ≃ *b*_*n*_*/c*_B_. In this linear response regime, *λ* diverges at the critical feedback strength. For 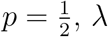 increases near the critical point as

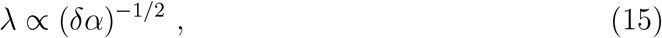

where *δα* = *α* − *α*^crit^. Because the Hill function *g*(*b*_*n*_) is non-linear, *λ* remains finite. The maximal value reached near *α*^crit^ scales with *c*_B_ as

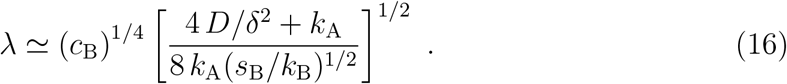

**Figure 3.**
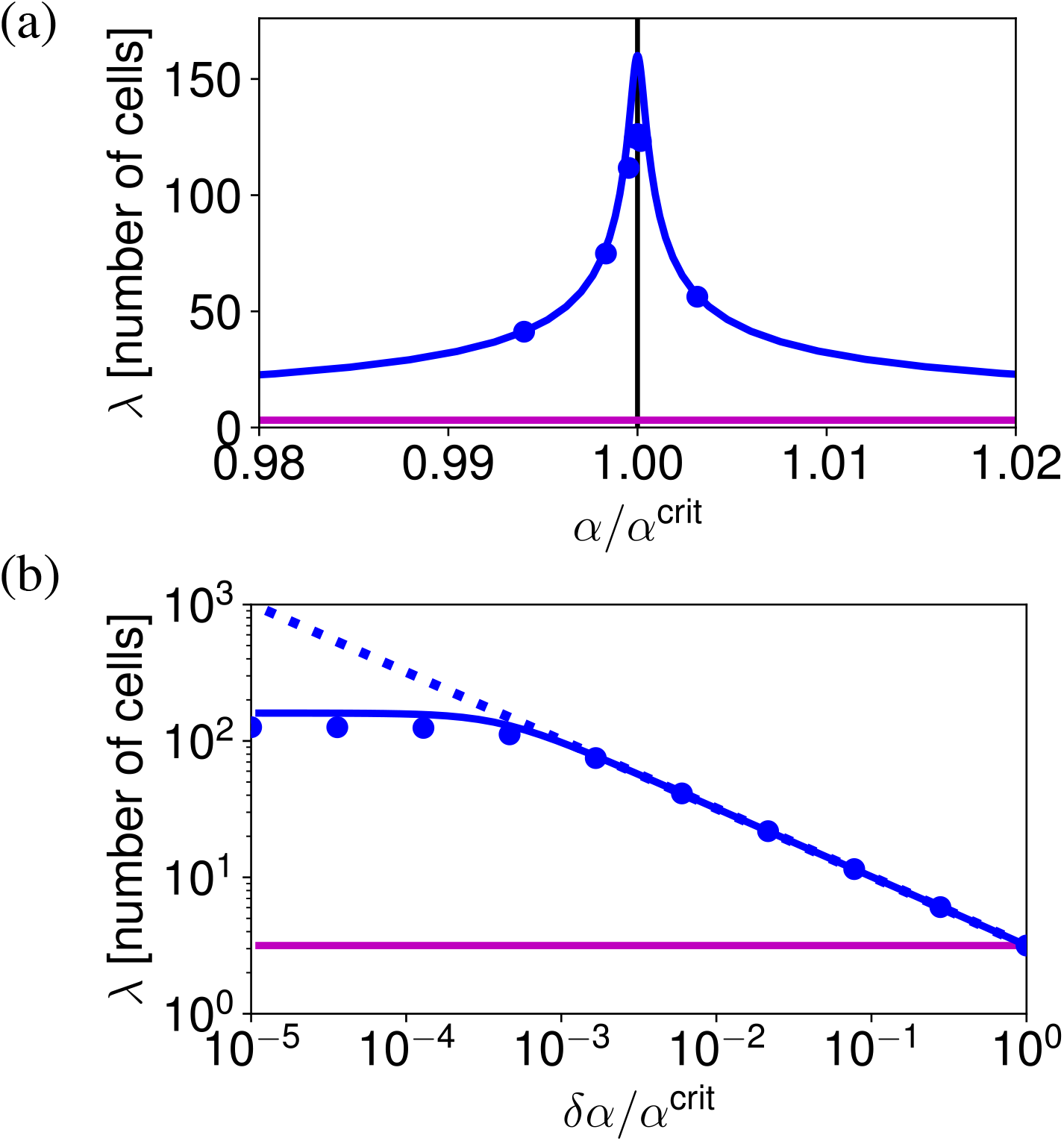
The decay length of the relay model reaches a maximum at the critical feedback strength *α*^crit^. (a) Decay length *λ* in cells as a function of the relative feedback strength *α/α*^crit^. (b) Decay length *λ* in cells as a function of the relative distance from the critical feedback strength *δα/α*^crit^, where *δα* = *α*^crit^ − *α*. The log-log plot exposes the scaling relationship stated in equation (15). Blue line: decay length according to the analytical approximation of steady-state solution, blue dots: decay length based on the numerical steady-state solution (see Appendix F), purple line: decay length of diffusion/degradation system, dashed blue line: decay length in the limit of a linear feedback response, vertical black line: *α* = *α*^crit^.

In the presence of secretion polarity oriented towards the distal end, 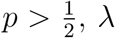 diverges near the critical point as

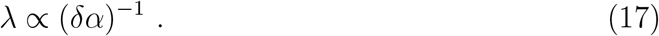

It is capped by the non-linearities at the maximal value given by

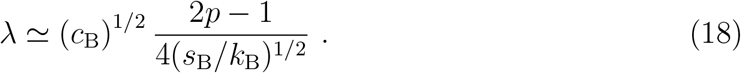

The maximal *λ* corresponds to a minimal value of *k*_eff_ which never reaches 0. Correspondingly, as a function of *α, λ* passes through a maximum near *α*^crit^, see figure 3(a), equations (16), (18).

Taken together, the positive feedback in the relay increases the decay length of the steady-state signaling gradient compared to diffusion and degradation, see figure 3. Moreover, the range of the signaling gradient can be regulated solely by changes in the feedback strength *α*. This is different from gradient formation by diffusion and degradation alone, where the decay length of the gradient is regulated by changes of the degradation rate or the diffusion coefficient. We next focus on the impact of secretion polarity on gradient range.

### 3.4. The influence of the emergent morphogen drift on gradient range

In the signaling relay, an emergent drift with effective velocity *v*_eff_ arises as a consequence of polarized secretion of morphogen, with 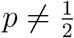, see equation (11). Thus, the emergent morphogen drift *v*_eff_ occurs in the absence of an actual directed transport of molecules through the system. In particular, the emergent drift has positive values for secretion polarity oriented towards the distal end i.e. for 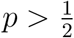. We observe that in this case the emergent drift increases the decay length of the steady-state signaling gradient reached at the critical feedback strength *α*^crit^, see figure 4, compare equations (16) and (18). This dependence of decay length on effective drift is also captured by the continuum description, see equation (8).

**Figure 4.**
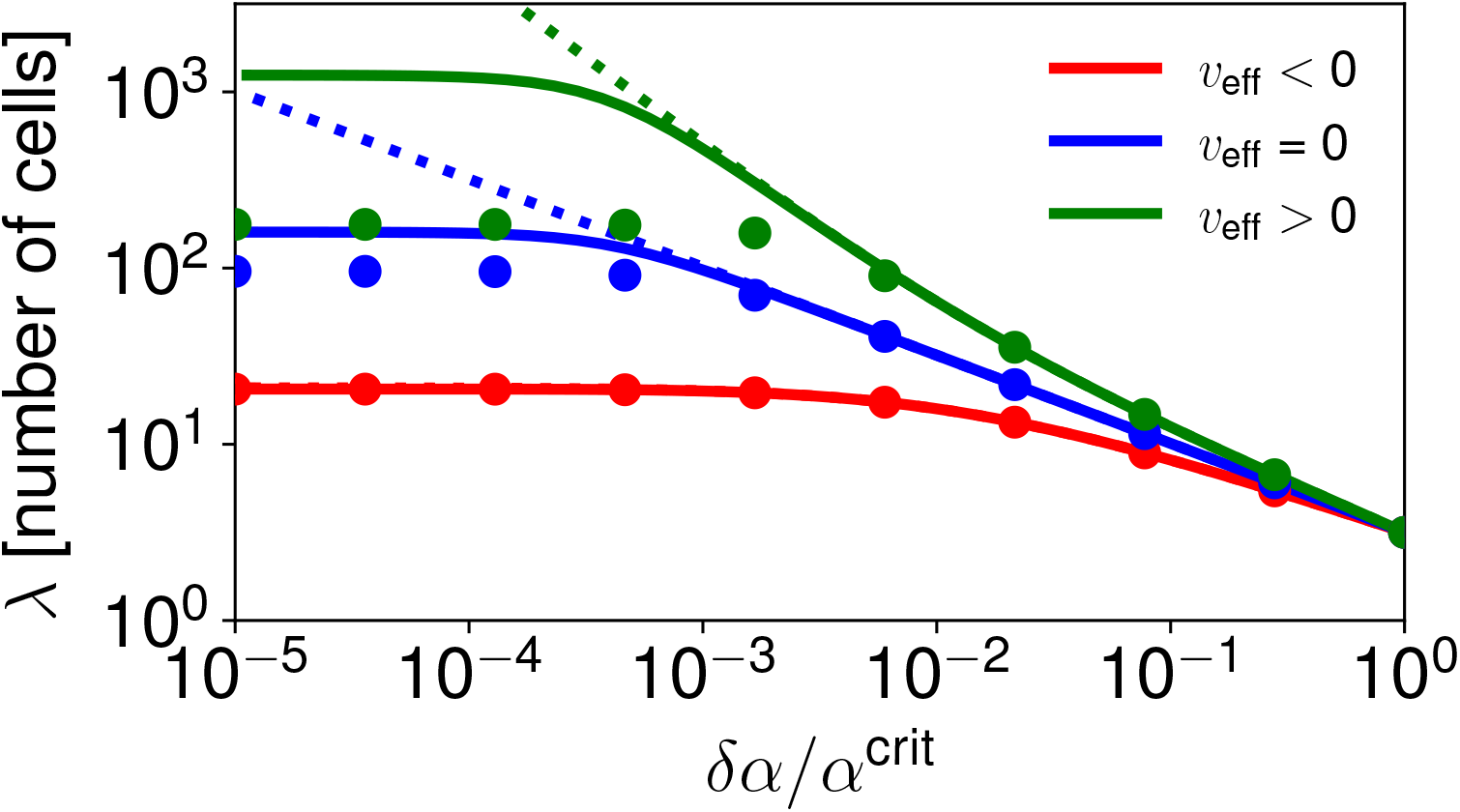
Polarized secretion towards the distal end (*v*_eff_ *>* 0) increases the decay length of the signaling gradient at steady state. Solid lines: decay length according to equation (13), dots: decay length based on the numerical steady-state solution of equations (1) and (2), dashed lines: decay length in the limit of a linear feedback response valid for signaling activities *b*_*n*_ much smaller than the Hill activation threshold *c*_B_. Blue: absence of secretion polarity, *p* = 0.5, green: preferred secretion towards the distal end, *p* = 0.99, red: preferred secretion towards the proximal end, *p* = 0.01.

The emergent morphogen drift also influences the sensitivity of the system to changes in the feedback strength as the feedback approaches its critical strength, compare equations (15) and (17). Thus, for preferred secretion towards the distal end, the decay length increases more rapidly as the feedback strength approaches its critical value as compared to the case without effective drift 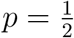, see figure 4.

Taken together, polarized secretion can increase the decay length of the steady-state gradients and can make the decay length more responsive to changes in the feedback strength. We next investigate how the feedback strength influences the dynamics of the system, in particular the time needed to build the gradient.

## 4. Dynamics of gradient formation with cell-to-cell signaling relay

In order to exert a patterning function, a morphogen gradient has to form. Therefore, apart from the decay length of the steady-state profile, the time it takes to form a profile is an important property of gradient formation. We use the slowest relaxation time of the system to reach steady state as a measure of the time it takes to form a gradient. We define the slowest relaxation time as the slowest exponential relaxation of concentrations close to steady state

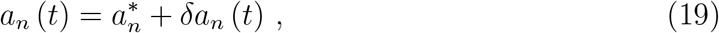

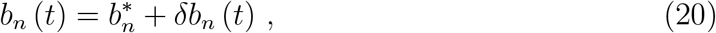

where *a*_*n*_ (*t*) and *b*_*n*_ (*t*) denote the concentration profiles of the morphogen and the intracellular signaling molecule at time 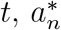 and 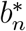 denote the respective steady-state concentration profiles and *δa*_*n*_ (*t*) and *δb*_*n*_ (*t*) the respective time-dependent deviations from it. Using the numerically determined steady-state profiles, we can then obtain the relaxation modes numerically, for details see Appendix G, see figure 5 circles. We observe that the relaxation time increases as *α* increases and that the dependence on *α* is steepest close to *α*^crit^, see figure 6. Compared to the case 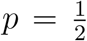, the relaxation time close to *α*^crit^ becomes shorter when secretion polarity is oriented towards the distal end, 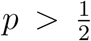, see figures 5 and 6. We compare these results to the relaxation modes obtained based on the approximation of the steady-state profile discussed in section 3.1, equation (6). This approximation works well away from the critical feedback strength but underestimates the relaxation time near *α*^crit^, figure 5 solid lines. Using a linear feedback, *g*_lin_(*b*_*n*_), provides a good approximation for the relaxation time away from the critical feedback strength, but leads to a divergence of the relaxation time at *α*^crit^, figure 5 dashed lines.

**Figure 5.**
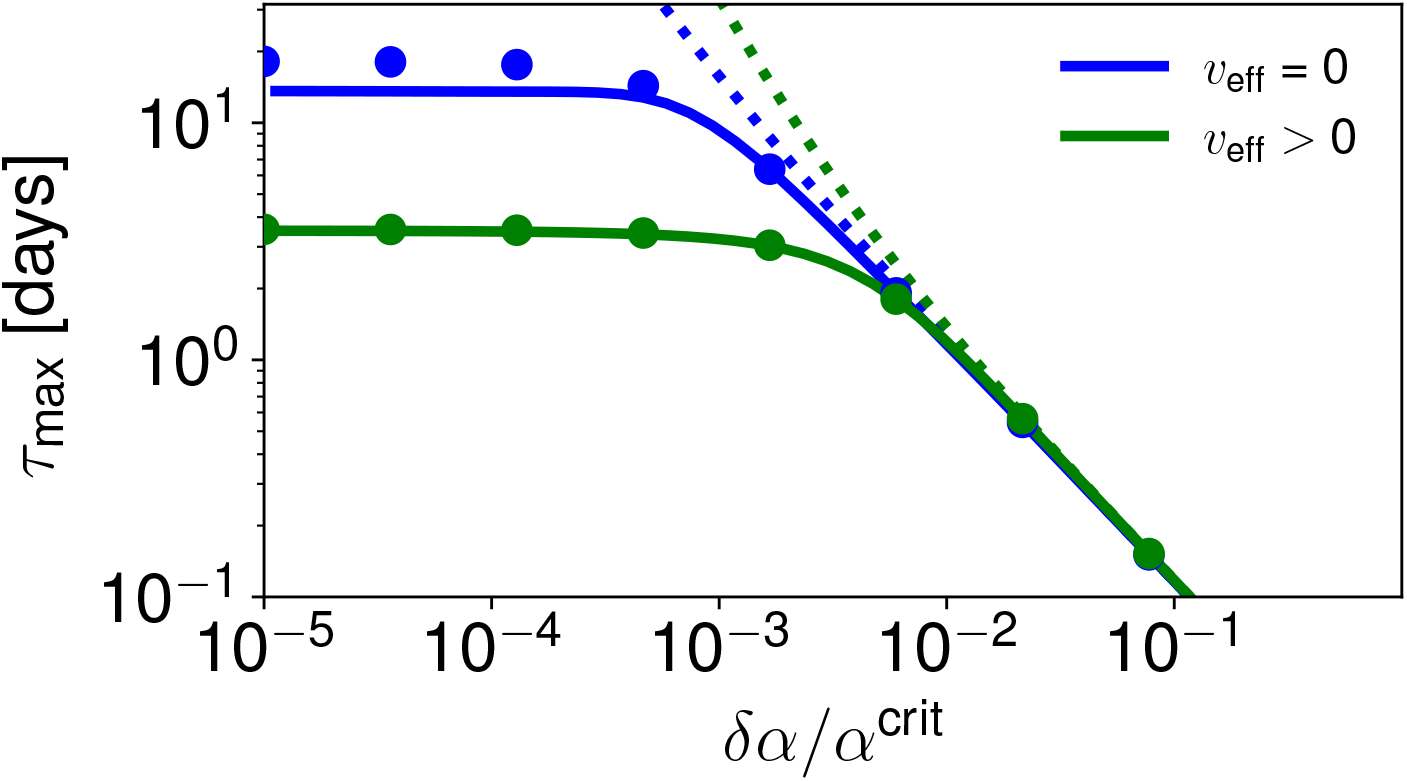
The relaxation time of the signaling-relay mechanism reaches a finite value at the critical feedback strength. The dots show the relaxation time obtained by linearizing the dynamics around the numerically obtained steady-state solution, the solid lines show the relaxation time obtained by linearizing the dynamics around the analytical approximation to the steady-state solution. The dotted lines show the relaxation time in the limit of a linear feedback response, i.e. in the absence of feedback saturation. The presence and direction of effective drift is indicated by the color code as specified in the legend: Blue: absence of secretion polarity, *p* = 0.5, green: preferred secretion towards the distal end, *p* = 0.99. *δα* = *α* − *α*^crit^.

**Figure 6.**
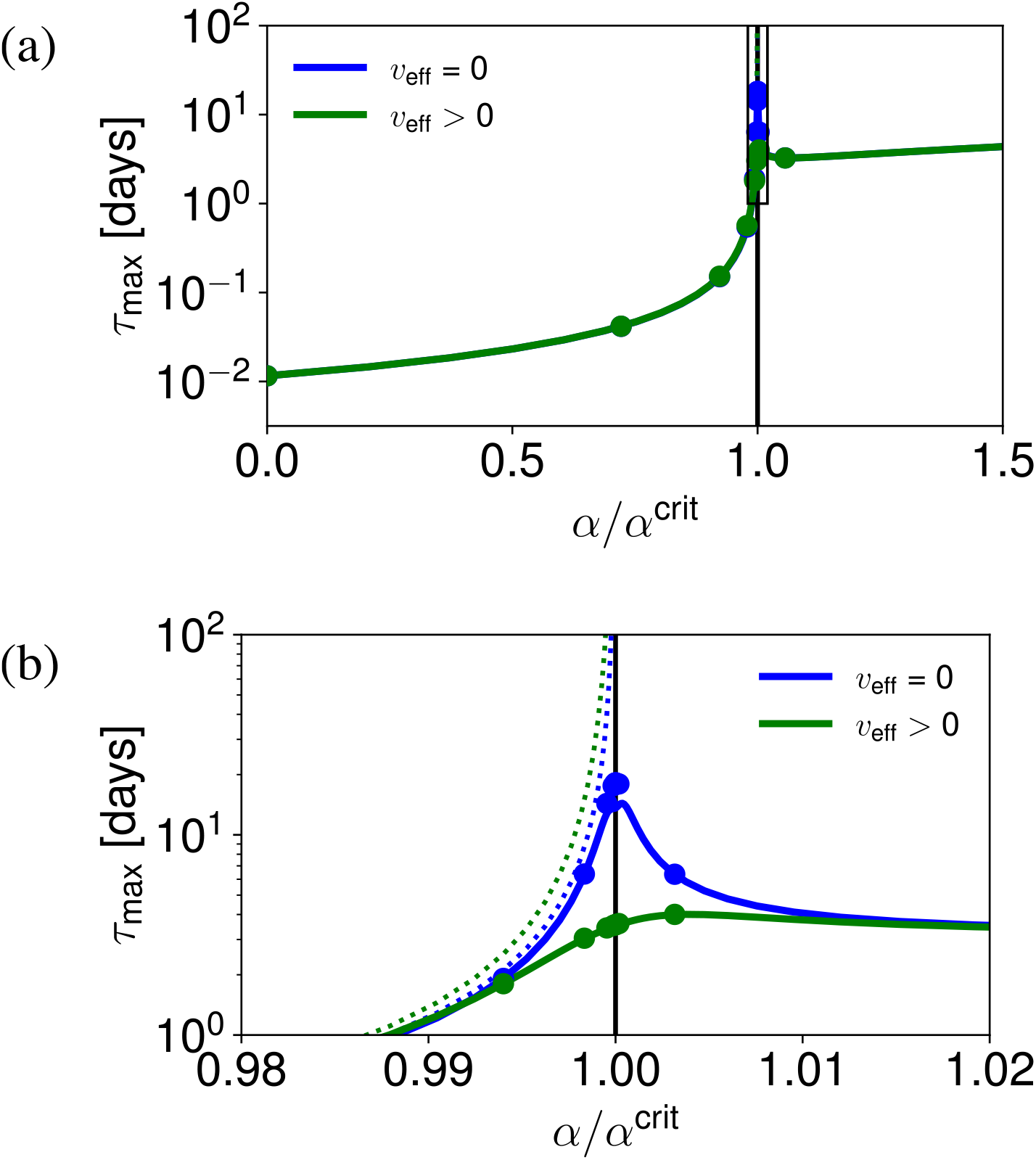
The relaxation time of the relay mechanism reaches a local maximum close to the critical feedback strength. (a) The relaxation time as a function of the feedback strength. (b) Blow-up of the boxed region in (a). Dots show the relaxation time obtained by linearizing the dynamics around the numerically obtained steady-state solution, solid lines show the relaxation time obtained by linearizing the dynamics around the analytical approximation to the steady-state solution, dotted lines show the relaxation time in the limit of a linear feedback response, i.e. in the absence of feedback saturation. The vertical black line marks the critical feedback strength. The presence and direction of drift is indicated by the color code as specified in the legend: Blue: absence of secretion polarity, *p* = 0.5, green: preferred secretion towards the distal end, *p* = 0.99. *δα* = *α* − *α*^crit^.

Taken together, the relaxation time slows down with increasing feedback strength *α* and reaches a local maximum near *α*^crit^ where it stays finite due to the non-linearities in the feedback, see figure 5. However, different from the decay length, the relaxation time is very asymmetrical around *α*^crit^, reaching higher levels for *α > α*^crit^ as compared to *α < α*^crit^. Secretion polarity towards the distal end can speed up the relaxation times close to *α*^crit^, see figures 5 and 6.

## 5. Scaling of decay length and relaxation time in gradient formation

Patterning of tissues requires a range of gradient decay lengths from tens of micrometers up to of the order of millimeters. In order to exert a patterning function, those gradients have to be formed on time scales compatible with the respective patterning process, for instance days to weeks in case of planarian regeneration. In this section, we discuss the scaling of relaxation time with decay length as the decay length is increased by changing the feedback strength.

As discussed above, the decay length of the gradients at steady state is largest for feedback strengths close to the critical one at which it reaches a maximum, see figure 3(a). In contrast, the relaxation time reaches a local maximum close to *α*^crit^ and stays high at values *α > α*^crit^, see figure 6. Therefore, generating gradients is fastest at feedback strengths below and up to the critical feedback strength, see figure 7. Figure 7 shows the scaling of relaxation time *τ*_max_ with decay length *λ*. We observe that for 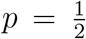, i.e. vanishing effective drift *v*_eff_, the relaxation time scales as *τ*_max_ ∞ *λ*^*ν*^ for *α < α*^crit^, with *ν* ≃ 2, see figure 7. In other words, doubling the decay length increases the relaxation time by a factor of four. This is the same scaling relationship observed for gradient formation by diffusion and degradation, see Appendix A, equation (A.8). Note that for gradient formation by diffusion and degradation, the decay length and the relaxation time are regulated by changes in the degradation rate, whereas in the signaling relay discussed here, they are regulated by changes in the feedback strength *α*.

**Figure 7.**
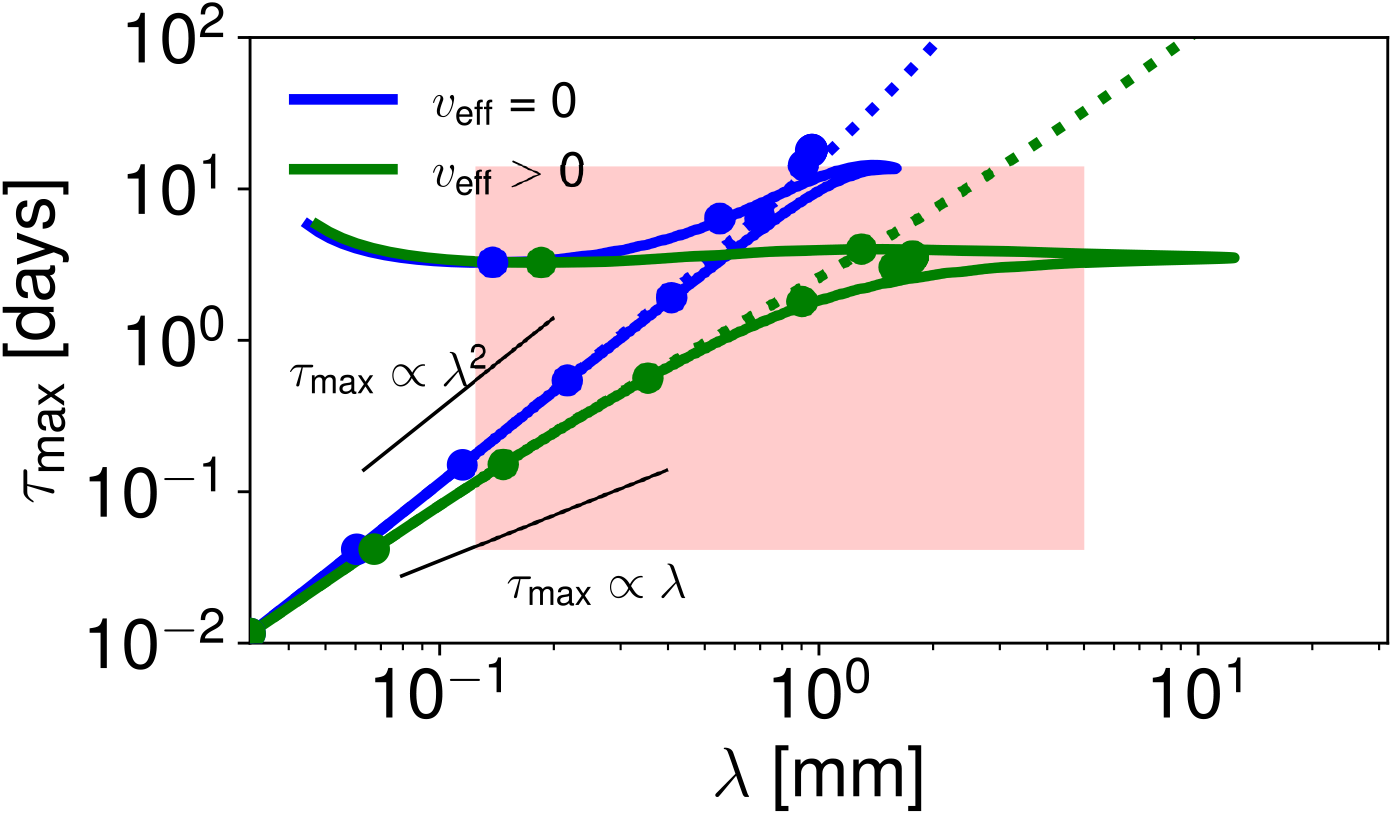
The trade-off between the decay length and the relaxation time of the relay model for changing feedback strength. The dots show the relaxation time based on dynamics linearized around the numerical steady-state solution and the decay length based on the numerical steady-state solution. The solid lines show the relaxation time based on the dynamics linearized around the approximate analytical steady-state solution and the decay length according to equation (13). The dashed lines show the relaxation time and the decay length in the limit of a linear feedback response, i.e. in the absence of feedback saturation. The presence and direction of drift is indicated by the color code as specified in the legend: blue: absence of secretion polarity, *p* = 0.5, green: preferred secretion towards the distal end, *p* = 0.99. The black lines above and below the curves show a scaling behavior of *τ*_max_ ∞ *λ*^2^ and *τ*_max_ ∞ *λ*, respectively. *τ*_max_ ∞ *λ*^2^ is the scaling behavior of gradient formation purely by diffusion and degradation. The region shaded in red corresponds to length and time scales relevant for planarian regeneration, i.e. 0.125 mm to 0.5 cm and 1 h to 14 days, respectively.

For secretion polarity oriented distally, 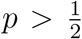, the scaling relationship is altered, with *ν <* 2, see figure 7. As *λ* reaches a maximum, deviations from these scaling relations occur. These are most pronounced when estimated using our approximations, see solid lines in figure 7. Taken together, long-ranged steady-state profiles are reached faster when formed by a molecular relay including secretion polarity compared to diffusion and degradation alone.

## 6. Discussion

In this paper, we examined the potential of a cell-to-cell signaling relay in morphogen gradient formation. In addition to diffusion and degradation, our model accounts for positive feedback of the morphogen on its own production across the entire cell field for gradient formation. This morphogen-mediated morphogen production is generated by a positive feedback loop, consisting of two elements. Firstly, morphogen production is positively regulated by intracellular morphogen-induced signaling activity and secondly, this intracellular signaling activity is positively regulated by the extracellular morphogen concentration, see figure 1. This way, the signal is propagated between neighboring cells, as the whole cell field becomes a signaling-dependent source. The overall strength of the feedback is spatially homogeneous across the cell field and set by the parameter *α*. Gradient initiation and orientation are set by a local, signaling-independent morphogen source at the proximal boundary. The non-linearity of the feedback response, in particular its saturation behavior, ensures that the system is not filled up with signaling molecules. Instead, the non-linear feedback combined with the linear degradation of the morphogen generates morphogen and signaling gradients emanating from the local, signaling-independent source. In conclusion, different from gradient formation by diffusion and degradation alone, the cell-to-cell relay provides an additional route of signal propagation. This propagation is due to the fact that morphogen signaling feedbacks on its own production, thus turning the entire cell field into a signaling-dependent source and this way having substantial influence on pattern formation.

We are not the first to consider the potential of a signaling relay in gradient formation. When putting forward their bucket brigade model of gradient formation, in which morphogens travel between receptors to form a receptor-bound gradient, Kerszberg and Wolpert briefly discussed whether such a bucket brigade could be replaced with an “induction cascade”, i.e. a positive feedback loop [45] similar to the one experimentally observed in *Xenopus* development by Reilly et al. [48]. Kerszberg and Wolpert postulated that gradient formation by relay required precise signal modulation as the signal was passed on from one cell to the next leading to the production of slightly less morphogen than sensed [45]. They state that the versions of positive feedback they have tried either propagated the signal indefinitely or let to the signal dying out too quickly [45]. In contrast to these considerations, we here show a possible realization of a cell-to-cell relay accomplished by a positive feedback loop that leads to the formation of long-ranged morphogen and signaling gradients. Importantly, the signaling relay in our model is realized as a feedback loop, that is, the extracellular morphogen concentration and the intracellular signaling activity regulate each other. This is a self-organized system that can generate states in which the morphogen production is spatially modulated and the morphogen concentration is in parallel spatially modulated. This can be realized for homogeneous feedback strengths *α* across the whole cell field because the feedback of signaling activity on morphogen production is modulated by the local signaling activities, see equation (3). The latter firstly ensures a response to the local levels of signaling activity and secondly provides feedback saturation for increased signaling levels. Together with the degradation of morphogen, see equation (1), this allows for a steady state and prevents runaway morphogen production despite the positive feedback. In the absence of feedback saturation, there exists a critical point in the feedback strength at which production due to the positive feedback would exceed the degradation and no steady-state would be reached. In the presence of feedback saturation, the system still reaches a steady state at this critical feedback strength. However, for feedback strengths above the critical one, gradient formation is inefficient, see figure 6, and gradients become rather flat, see figures D1, D2 in the appendix. At the critical feedback strength the decay length of the gradient profile becomes maximal, see figure 3. For even larger feedback strengths, the gradient decay length decreases again, see figure 3, and profiles become rather flat with a high base level of morphogen due to the positive feedback and a shallow spatial profile, see figures D1, D2 in the appendix. We speculate that in this regime, the signaling relay may lead to traveling fronts, similar to those reported in [49]. Overall, our results demonstrate that the formation of steady-state gradients is indeed possible in systems incorporating a signaling relay and that the gradient decay length depends on the feedback strength.

One interesting feature of the inclusion of a signaling relay into morphogen gradient formation is that it affords the extension of gradient decay lengths beyond the decay length set by diffusion and degradation alone. We make a conservative choice of the molecular diffusion coefficient of 1 *µ*m^2^*/*s to take into account the slow diffusivity of lipid-modified molecules compared to e.g. Fgf8 (*D* = 53 ± 8 *µ*m^2^*/*s [16]), and use a degradation rate of 10^*−*3^ s^*−*1^, like the one measured for Wg [15] or Fgf8 [16]. With these numbers, we show that the signaling relay can generate gradients with decay lengths of up to about 950 *µ*m within a time scale of days, see figures 4, 6. When the signaling relay is combined with secretion polarity towards the distal end, i.e. preferred secretion on the cell boundary facing away from the local, signaling-independent source, 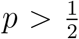, it can reach decay lengths of up to 1770 *µ*m in this scenario, see figure 4. In contrast, gradients formed by diffusion and degradation alone would reach a decay length of 31 *µ*m for the same diffusion coefficient and degradation rate. Thus, the signaling relay leads to an increased range of the gradient compared to a gradient formed by diffusion and degradation alone, see figure 3. The decay length-extension effect is due to the following components: First, production of morphogen throughout the tissue due to the signaling relay gives rise to morphogen profiles that reach large distances even though individual molecules only travel shorter distances. The morphogen profile still follows a transport equation (equation (8)) which is characterized by effective transport coefficients (equations (9), (11)) and an effective degradation rate (equation (10)). The effective transport coefficients describe the effective diffusion coefficient and drift velocity associated with net transport even though individual molecules do not drift and exhibit only limited molecular diffusion. The increased effective diffusion (equation (9)) already leads to an increase in the decay length of the gradients. Furthermore, the signaling relay also leads to an effective degradation of the molecule that is slower than the molecular degradation (equation (10)) because molecules are newly produced across the whole cell field. The joint effect of an increased effective diffusion coefficient and decreased effective degradation rate is that the resulting signaling pattern can extend beyond the limits set by the molecular diffusion coefficient and degradation rate of the morphogen. Taken to its extreme, this means that with the signaling relay, morphogen gradients can form even in the limit of no molecular diffusion, exclusively due to the positive feedback and the cell-to-cell relay. This effect of an increased decay length can be further enhanced by the polarized release of the morphogen on the cell boundary facing away from the local, signaling-independent source, 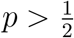. This could be mediated by a biological mechanism like planar cell polarity [50]. Polarized secretion of the molecules produced in response to the positive feedback leads to the emergence of an effective drift of morphogens through the system (equation (11)) even though individual molecules do not exhibit drift. This further increases the decay length of the gradients formed, see figure 4. Thus, secretion polarity is an additional handle to increase the range of gradients formed by a signaling relay. Overall, a signaling relay allows to reach decay lengths of the order to millimeters even for slowly diffusing molecules, that is one to two orders of magnitude larger than what is achieved by diffusion and degradation alone.

A further feature of the signaling relay is that it also provides an interesting opportunity to regulate the decay length of the morphogen and signaling gradients by tuning the feedback strength. The signaling relay modulates the effective transport coefficients and the effective degradation rate that together determine the decay length of the gradient. The signaling relay hence provides a new way of regulating gradient decay length compared to diffusion-degradation gradients. For diffusion-degradation gradients, the range of profiles is regulated by tuning the degradation rate or the diffusion coefficient of the morphogen. The presence of a critical point in the signaling relay provides a sensitive response to the feedback strength. This can allow the regulation of gradient decay lengths by small changes to the signaling feedback strength close to the critical feedback strength, see figure 3. Such changes could be achieved by altering the amount of morphogen produced in response to the same signaling level, e.g. by modulating the signal amplification of the signaling pathway. Such a regulation of gradient decay length via signaling provides a new scenario for how gradient scaling could be achieved [51]. For example, it will be interesting to further investigate how the feedback strength could be self-tuned close to the critical feedback strength such that the decay length follows the organism size. In conclusion, the signaling relay harbors attractive concepts to explore within the scope of the unsolved problem of pattern scaling.

Apart from the decay length of a morphogen gradient, the time it takes to form the profile is an important characteristic of any patterning mechanism, since the profile has to form on time scales relevant for the pattering process in question. Some morphogen gradients exert a pattering function already before they have reached steady state [52]. However, the relaxation time of a morphogen gradient, defined as the slowest exponential relaxation time close to the steady state, is a good measure of the dynamics of the gradient. In general, the formation of more long-ranged gradients takes longer compared to the formation of more short ranged profiles. Accordingly, we find that for a cell-to-cell signaling relay for *α < α*^crit^, the relaxation time slows down with increasing feedback strength *α*, see figure 5, as the decay length increases, see figure 3. As *α* increases beyond the critical feedback strength, the decay length decreases again, see figure 3, while the relaxation time stays high for *α > α*^crit^, see figure 6. However, secretion polarity towards the distal end can speed up the relaxation times in particular near *α*^crit^, see figures 5, 6, while increasing the decay lengths, see figure 4.

To quantify this trade-off between the decay length and the relaxation time, we analyzed the scaling between the decay length and the relaxation time, *τ*_max_ ∞ *λ*^*ν*^. In the signaling relay-model, the relaxation time scales with the decay length with an exponent of *ν* ≃ 2 in the absence of secretion polarity for *α < α*^crit^, see figure 7. This is the same scaling relationship observed for gradient formation by diffusion and degradation. However, in the presence of secretion polarity towards the distal end, this scaling exponent is smaller than two for *α < α*^crit^, see figure 7. Thus, positive feedback combined with polarized secretion allows the formation of long-ranged morphogen gradients in a shorter time compared to gradient formation by diffusion and degradation. Since the change in decay length is achieved by changing the feedback strength rather than the degradation rate, long-ranged signaling gradients can be formed for measured values of the degradation rate and the diffusion coefficient. In particular, it is not necessary to assume particularly long-lived or very quickly diffusing morphogens. In contrast, it is necessary to assume high turnover, i.e. short life-time, of the intracellular signaling molecules as we did in figures 4 - 7, *k*_B_ ≪ *k*_A_. In general, it is more likely that cells have the ability to decrease molecular life times by up-regulating the degradation machinery and/or preferentially marking the molecules in question for degradation than to make molecules more long-lived. Thus, postulating high turn-over rates of the intracellular signaling molecule is closer to biological reality than postulating very long life times for the morphogens. Taken together, the relay allows the formation of millimeter-ranged signaling gradients on biologically relevant time scales of hours to days for a physiologically relevant choice of parameters.

Gradient formation by a signaling relay as discussed in this paper results in morphogen-production gradients emanating from the signaling-independent source. In contrast, in gradient formation by diffusion and degradation, production is typically thought to be restricted to a local source at the boundary of the system, similar to the signaling independent local source at the proximal boundary of the system discussed in this paper. Thus, morphogen expression gradients may serve as an experimentally accessible signature of a signaling relay. Once the potential for a signaling relay has been identified by the presence of such expression gradients, the mutual dependence of the morphogen and the signaling gradient can be assayed to experimentally identify a signaling relay: While the intracellular signaling levels are strictly downstream of a classical morphogen gradient, they are instrumental for the formation of the morphogen gradient in a signaling relay. Perturbation of the signaling levels should hence not affect the morphogen gradient if it were formed by diffusion and degradation. In contrast, such perturbation should disrupt the formation of a morphogen gradient formed by a signaling relay. In planarians, expression gradients of Wnt are observed [32, 46], suggesting the presence of a signaling relay. In order to test the interdependence of the morphogen (Wnt) and the signaling activity (*β*-Catenin) gradient, it will be necessary to measure the two independently.

There is experimental evidence for the functional importance of signaling relays in other biological systems beyond of the Wnt signaling gradient organizing the main body axis in planarians [32]. For instance, signaling relays seem to be required for mesoderm induction during early embryonic development of *Xenopus* [48]. Further, *wnt* expression gradients emanating from the hypostome are observed in Hydra [53]. Additionally, there is Wnt expression restricted to the tip of the hypostome [54] that might be related to the signaling-independent source. It will thus be interesting to investigate whether a signaling relay of the kind discussed in this paper is involved in patterning the main body axis in Hydra that is known to be organized by Wnt signaling [54].

Moreover, it will be interesting to revisit cases in which patterning is believed to occur by classical morphogen gradients and to analyze if in fact there is a signaling relay at play. For instance, it could be examined whether the shape of the Wg gradient in the *Drosophila* fly wing disk changes in the absence of the *Drosophila β*-Catenin homologue Amardillo; or whether the Dpp gradient still forms correctly in the absence of Smad. Interdependencies between the signaling and the morphogen gradient in these two well-known examples of gradient formation could point towards a signaling relay being present in these systems. Further, the main *Drosophila* Wnt, Wg, might be subject to a signaling relay by a positive feedback loop like discussed in this paper as it has a TCF binding site in its regulatory region [55]. It could thus be regulated by the intracellular signaling levels manifested as Amardillo (the *Drosophila β*-Catenin homologue) that binds to TCF activating expression of genes with TCF binding sites. In fact, there is evidence for positive feedback in *wg* expression in intersegmental patterning during *Drosophila* development [56]. In light of these findings, it might be interesting to revisit the finding that membrane-tethered Wg does not impair pattering in the developing *Drosophila* wing imaginal disk [39]. Could it be that there is a signaling relay at play generating a Wg gradient with concentrations levels that so far have eluded detection?

Pattering by morphogen gradients has mostly been studied in a very few canonical model systems in molecular detail, specifically, in *Drosophila* embryos [15] and zebrafish embryos [16]. In these systems, patterning occurs on small length scales and is well explained by diffusion and degradation [15, 16]. However, patterning is not restricted to embryonic development with its small structures. In contrast, it is also required on adult length scales, notably during regeneration. Research in planarians flatworms has suggested that long-ranged gradients might be formed by a signaling relay [32]. In this paper, we thus theoretically investigated the potential of signaling relays for the formation of morphogen gradients, in particular on long length scales. In a signaling relay, morphogens do not have to propagate through the whole system by diffusion. Instead, they are produced within the target tissue itself, that becomes a signaling-dependent source thanks to positive feedback. Signal propagation by positive feedback in a signaling relay is thus an attractive concept for gradient formation in contexts in which it is not feasible to propagate the signal by diffusion into the target tissue, either because of the size of the system or because of the molecular properties of the morphogen or both in the case of the Wnt signaling gradient in planarians. It will be interesting to investigate the presence of signaling relays in different species, in particular those that are capable of regeneration and thus require pattering on large length scales, such as planarians, hydra and axolotl. Regeneration necessitates robust pattern formation. The self-organized nature of the signaling relay makes it a promising candidate for robust pattern formation. Thus, signaling relay is an attractive concept for long-range, robust pattern formation, as required for instance during regeneration. Further, it allows rationalizing the formation of long-ranged gradients by slowly diffusing morphogens such as Wnt and Hhg.

## Acknowledgments

We thank Jonathan Bauermann, Charlie Duclut, Steffen Werner, Daniel Aguilar-Hidalgo, Alexander Mietke, Felix Meigel, Hanh Vu, Mario Ivanković and James Cleland for insightful discussions.

## Appendix

### Appendix A. Continuum theory of signaling gradient formation

In this section, we recapitulate the continuum theory of gradient formation including drift. The diffusion/degradation model for gradient formation that has successfully been applied to explain the Dpp gradient in the wing imaginal disc [15] is contained in this model as the special case of no-drift. In particular, the dynamics of morphogen concentration *C* are given by secretion in a local source of width *S* with rate *s*, diffusion with diffusion coefficient *D*, degradation with rate *k*, and drift or advective transport of molecule concentration with drift speed *v*,

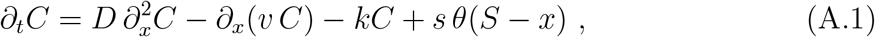

where *x* denotes the spatial coordinate in the dimension along which the gradient is formed, and *θ* denotes the Heavyside function, where *θ*(*y*) = 0 for *y <* 0 and *θ*(*y*) = 1 for *y* ≥ 0. In equation (A.1), 0 *< x* ≤ *L* with a tissue of size L and the source at the proximal boundary. Positive values of *v* indicate transport in positive *x*-direction. Thus, for the profile decreasing in positive *x*-direction, *∂*_*x*_*C <* 0, the drift term leads to drift down the concentration gradient.

This dynamics leads to steady-state concentration profiles of the form

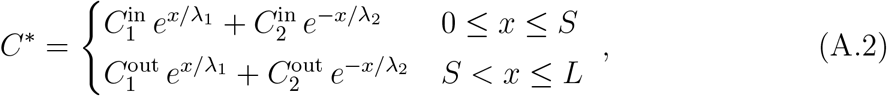

where the decay lengths are given by

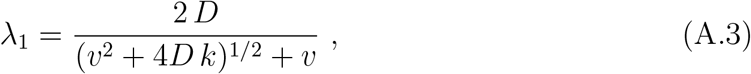

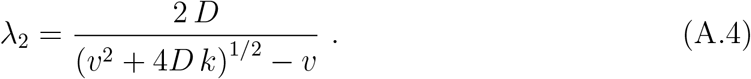

**Figure A1.**
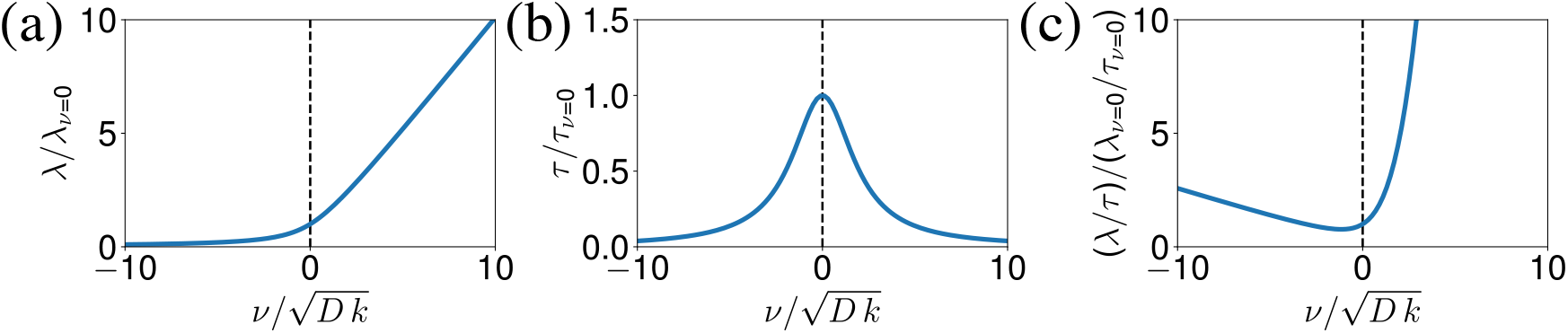
Comparison of the continuous diffusion/degradation model with and without drift. Comparison of steady-state profile decay lengths of the diffusion degradation model with drift (*λ*) to the one without drift (*λ*_*v*=0_) (a), of the respective relaxation times (b), and the trade-off between decay length and relaxation time (*λ/τ* and *λ*_*v*=0_*/τ*_*v*=0_, respectively (c). All plots as a function of a dimensionless parameter relating the influence of drift with speed *v* that is varied to the one of diffusion with diffusion coefficient *D* and degradation with degradation rate *k*, that are set to values measured in the fly wing disc (D = 0.1 *µ*m^2^, k = 10^*−*3^ s^*−*1^ [15]). The dashed lines indicate the absence of drift, *v* = 0.

Note that in the absence of drift, *v* = 0, this expression simplifies to the well-known decay length of a diffusion/degradation system that we denote by the index *v* = 0:

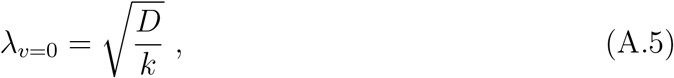

For no-flux boundary conditions, −*D ∂*_*x*_*C*(0) + *v C*(0) = −*D ∂*_*x*_*C*(*L*) + *v C*(*L*) = 0, and differentiability, i.e. matching value and flux at point *S*, where the source and the non-source regions meet, 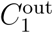 is very small and goes to zero as the system size *L* increases. Therefore, the profile outside of the source region is given by an exponentially decreasing profile with the decay length *λ* := *λ*_2_.

Comparing the decay lengths in the presence and absence of drift, we note that the decay length is increased in a system with drift for positive drift speeds, see figure A1(a). The relaxation time of gradient formation, defined as the relaxation time of the most slowly relaxing Eigenmode, that is a good measure of how long the system takes to reach steady state, is given by

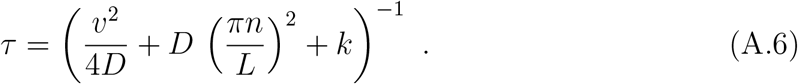

Again, the time-scale of the diffusion/degradation model is contained as the case *v* = 0:

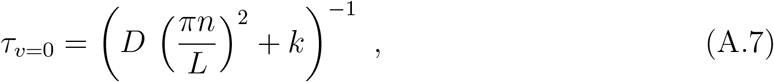

We can thus appreciate, that the system relaxes faster in the presence of drift, see figure A1(b) for a comparison in the limit of an infinitely large system (*L* → ∞).

There is a trade-off between the decay length of the steady state profile and the relaxation time of gradient formation. That is, the formation of long-ranged steady state profiles takes increasingly long. In particular, in the absence of drift, the relaxation time scales as

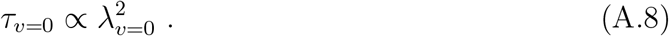

As we discussed above, in the presence of drift, the steady-state decay length in a system with positive drift speeds is increased compared to a system without drift (figure A1(a)), and the relaxation time of gradient formation is decreased (figure A1(b)). Thus, the trade-off between the decay length of the steady state profile and the relaxation time, that can be quantified as the ratio of the decay length and the relaxation time *λ/τ* is markedly improved, see figure A1(c). This trade-off is a measure of how long it takes to form a gradient of a given decay length. Thus, a gradient with an equal decay length is reached faster in the presence of drift or a gradient with a longer decay length is reached in the same time.

While drift of morphogen concentration through the system may very well explain the formation of signaling gradients on decay lengths in the millimeter range on biologically relevant time scales of hours to days, it leaves open the question of the origin of drift. Drift may arise due to concerted cilia beating or muscle contraction [44], but neither can explain the formation of long-ranged gradients in the context of regeneration in which the respective structures would have to be built first.

### Appendix B. Boundary conditions

We consider the case in which the system ends with an extracellular space 0 at the proximal end and an extracellular space *N* at the distal end, see figure 1. We thus need to specify the boundary conditions *∂*_*t*_ *a*_0_ and *∂*_*t*_ *a*_*N*_. We assume no diffusive flux at these boundaries. Moreover, we choose the volumes of the extracellular spaces at the system boundaries to be smaller than those in the bulk, in particular

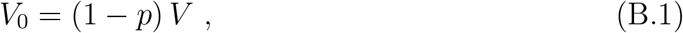

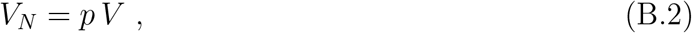

where *V* denotes the volume of the extracellular spaces in the bulk, i.e. *V*_*n*_ = *V* for *n* = 1, …, *N* − 1. This leads to the following dynamics at the system boundaries:

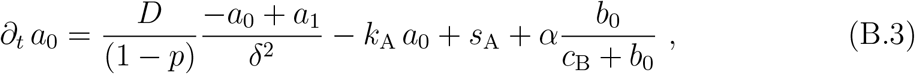

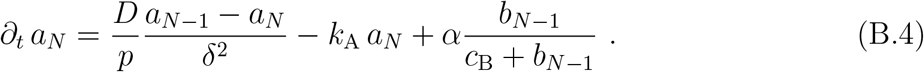

Note that with this choice of volumes at the boundaries, the secretion polarity in the production terms is balanced at the boundaries. To see this, consider that the cells produce individual molecules rather than concentrations and that thus the production rates measured in concentration per time depend on the volume of the extracellular spaces into which the produced molecules are secreted.

Note further that this choice of boundary conditions gives rise to flat profiles at the boundaries of the system, i.e. vanishing spatial derivative at the boundaries, see figure 2. Finally, note that in the presence of secretion polarity, 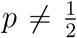, this choice of boundary conditions is not the same as no-flux boundary conditions.

### Appendix C. Numerical steady-state solution

We can obtain the steady-state solution to the signaling relay (equations (1), (2)) numerically by starting from an initial condition and computing the time evolution for long times. We choose the average of the first (*a*_0_ and *b*_0_, respectively) and the last value (*a*_*N*_ and *b*_*N−*1_, respectively) of the analytical approximation to the steady-state solution as initial conditions. In order to obtain the concentration profiles of the morphogen and the intracellular signaling molecule at time *t*, ***a***^*t*^, and ***b***^*t*^, respectively:

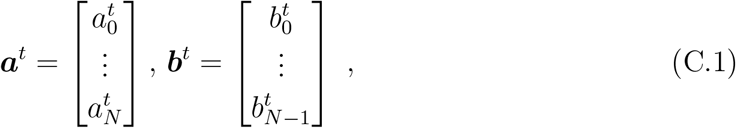

where the upper index refers to the time step and bold italic fond indicates vector notation, we need their concentration profiles at time *t* − 1, ***a***^*t−*1^, and ***b***^*t−*1^. We use a forward (explicit) Euler method to compute the (non-linear) dynamics of the concentration profile of the intracellular signaling molecule. Specifically, we add the change in concentration during one time-step of length Δ*t* to the current concentration profile ***b***^*t−*1^:

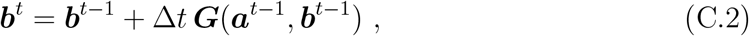

where the differential equation for the signaling activity ***G***(***a***^*t−*1^, ***b***^*t−*1^) is defined according to equation (2):

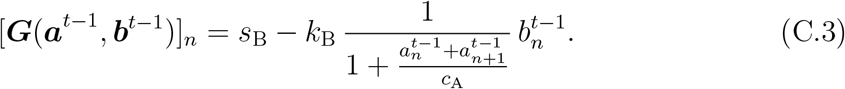

To compute the dynamics of the morphogen, we use an implicit-explicit Euler method [57]. In particular, we evaluate the linear parts (diffusion and degradation, described by the matrix ***N***) implicitly and the non-linear parts (i.e. the local, signaling-independent source, as well as the feedback, i.e. the signaling-dependent source, ***F*** (***b***^*t−*1^)) explicitly(see Ref. [57] for details on the method). This results in the following dynamics:

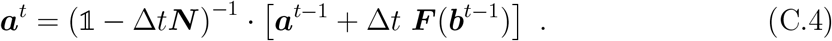

The linear part is given by diffusion and degradation:

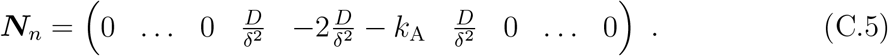

The non-linear part of the dynamics of the signaling molecule, ***F*** (***b***^*t−*1^), is given by the constant production in the local, signaling-independent source with rate *s*_A_ and the positive feedback according to equation (3). This results in:

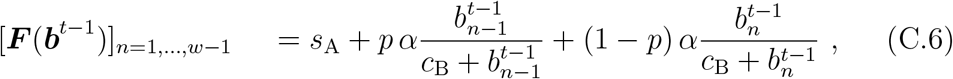

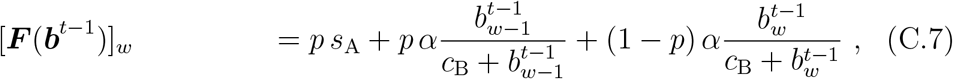

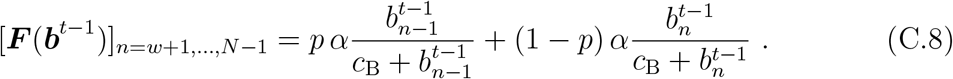

The boundary conditions are specified in the same way according to equations (B.3), (B.4). This way, we can compute the dynamic solution to our model and in particular, for long times, the steady-state solution. We verify that the steady-state is reached by observing that the profiles no longer change with time.

### Appendix D. Analytical approximation to the steady state

In short, we find an analytical approximation to the solution of the steady-state equation of the morphogen (equation (1) using equation (5)) by linearizing it around the piece-wise constant steady-state solution given by 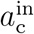 and 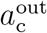 inside (*n* = 0, …, *w*) and outside (*n* = *w* + 1, …, *N*) of the source region, respectively. This piece-wise constant solution is defined by the constants solving the steady-state equations of the bulk of the source and the non-source region, respectively. We describe this procedure in more detail in this section:

We analyze the steady-state of the morphogen concentration in order to determine the piece-wise constant steady-state solution. Consider that at steady state, 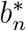 is linearly dependent on 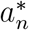 (equation (5)). Inserting this relationship (equation (5)) into the steady state equation for 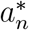 (equation (1)), we obtain

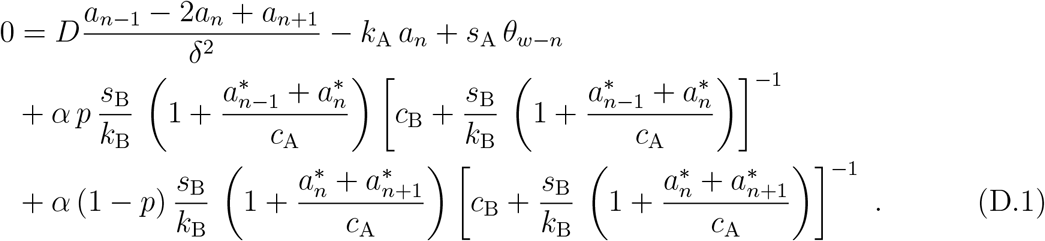

Based on this equation, we define 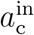 and 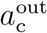 as the constants solving equation (D.1) for *n* = 1, …, *w* − 1 and *n* = *w* + 1, …*N* − 1, respectively. We obtain

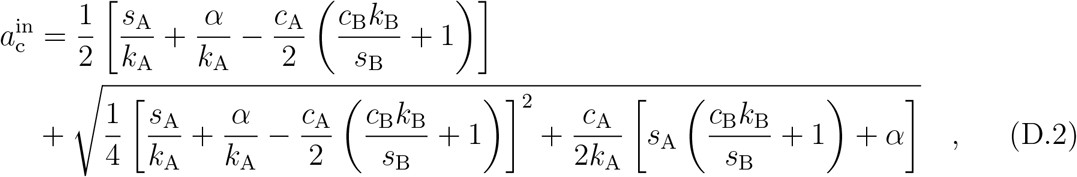

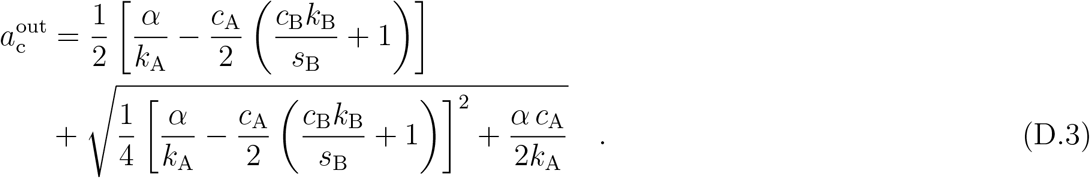

We then obtain the approximation to the steady-state solution by linearizing equation (D.1) around 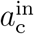 inside the source region for *n* = 0, …, *w*, and around 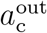 outside of the source region for *n* = *w* + 1, …, *N*, respectively. To this end, we express *a*_*n*_ as a deviation from these constants:

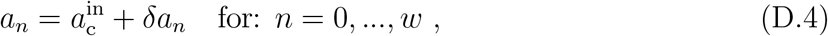

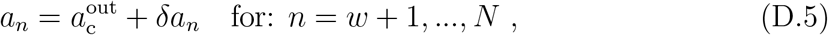

and linearize for small *δa*_*n*_. We thus obtain a set of linear equations:

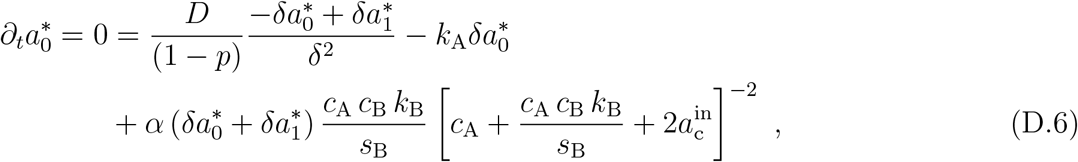

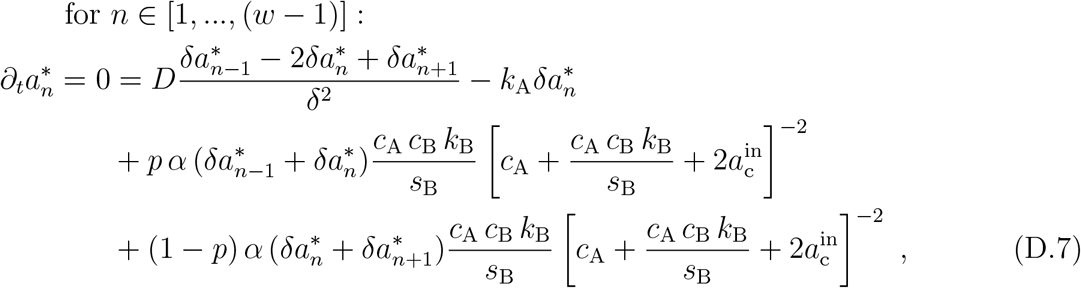

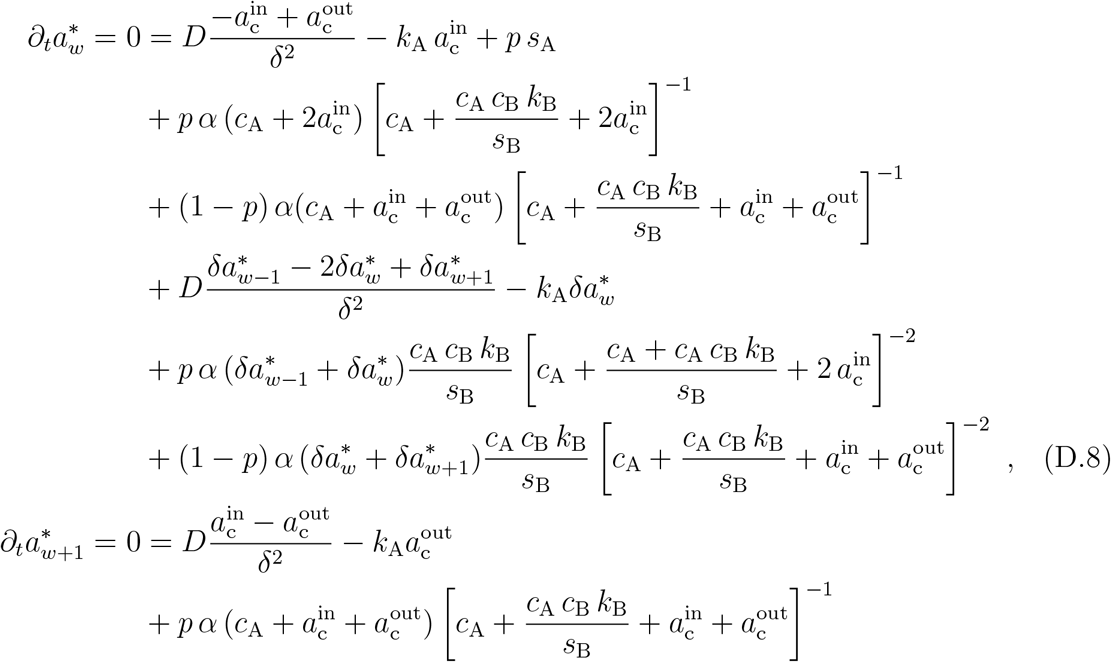

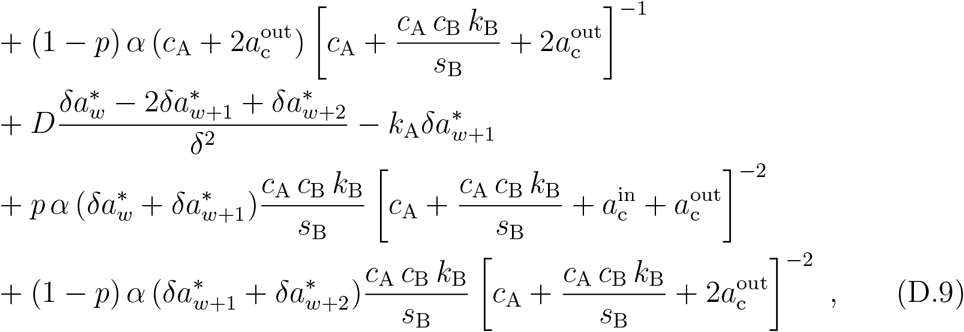

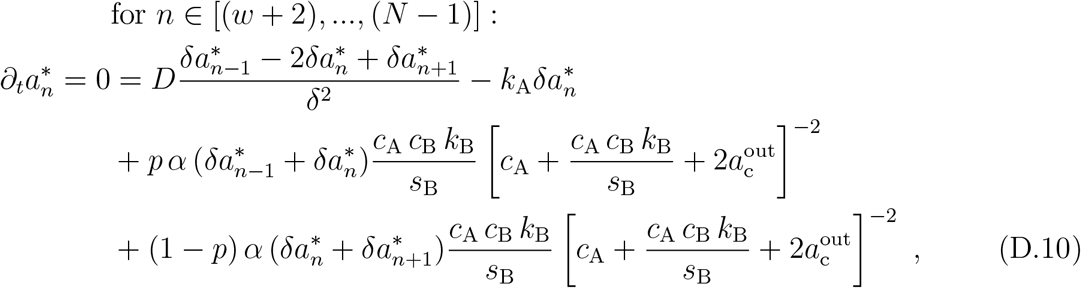

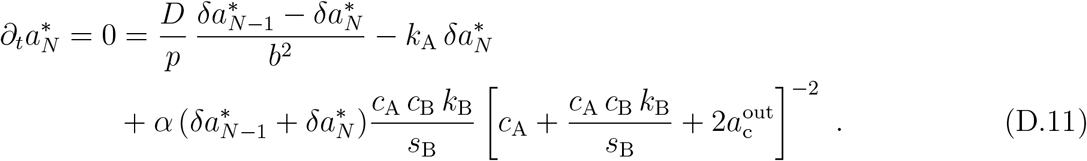

Note that the term for the signaling relay-dependent morphogen production by cell *w*at steady-state is given by 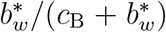. When re-writing this using equation (5), it contains 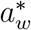and 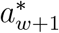. Therefore, it needs to be linearized around the sum of 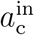 and 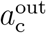. Thus, for 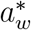 and 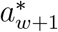, the constant term does not vanish (see equations D.8 and D.9) as it contains both 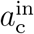 and 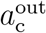.

We can solve these linearized steady-state equations using an exponential ansatz and obtain the solution given in equation (6). The constants in equation (6) are defined by the boundary conditions at *n* = 0 and *n* = *N*, as well as by the source/non-source interface at *n* = *w* and *n* = *w* + 1, equations (D.6), (D.8), (D.9), and (D.11). Plugging the exponential ansatz (6) into equation (D.6), (D.8), (D.9), and (D.11), we obtain:

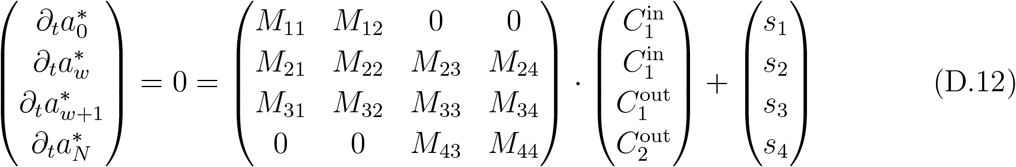

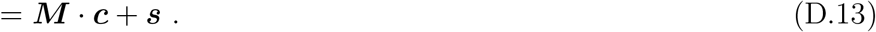

Thus, the constants in ***c*** are given as:

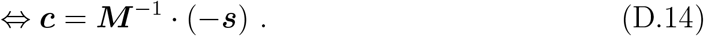

We obtain the decay lengths by expanding the exponentials in 1*/λ*. This expansion is valid for decay lengths much larger than one cell, i.e. *λ* ≫ 1, and thus the long decay lengths we are interested in. We find

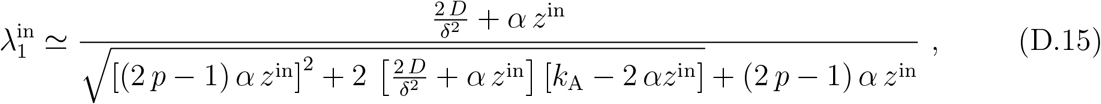

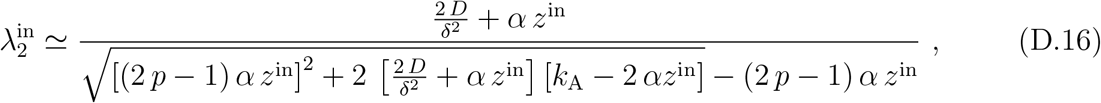

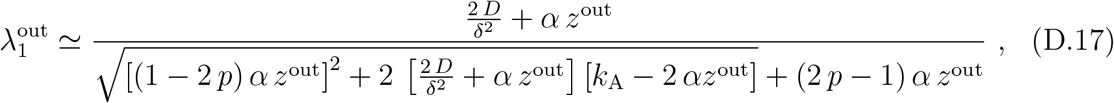

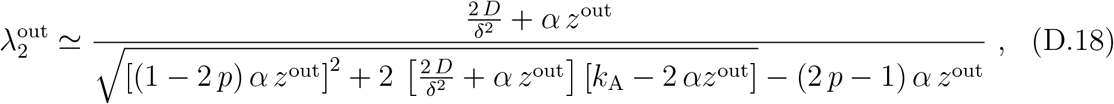

where

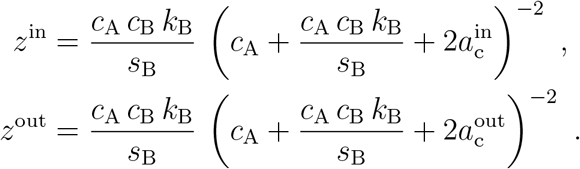

We can compare this approximation to the steady state to the steady-state profiles obtained numerically, see figures D1, D2. Note that the analytical approximation to the steady state is particularly good for feedback levels away from the critical feedback strength, see figures D1, D2. Note further that close to the critical feedback strength, the approximation is better for *b*_*n*_*/c*_B_ « 1, compare figure D1 (*c*_B_ = 1.66 × 10^8^ nM) and figure D2 (*c*_B_ = 1.66 × 10^3^ nM).

### Appendix E. Linearized dynamic equations

In order to calculate the effective transport coefficients and loss rate, defined by equation (8), we first determine eigenmodes of the dynamics linearized around the homogeneous steady-state solution given by 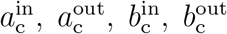, see Appendix D. The 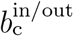 are obtained from the 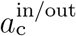 using equation (5). We discuss the dynamics outside of the source region. The dynamics inside of the source region can be obtained in the same way using the respective constants 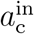 and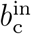. Outside of the source region, the dynamics linearized around the homogeneous steady state solution is given by

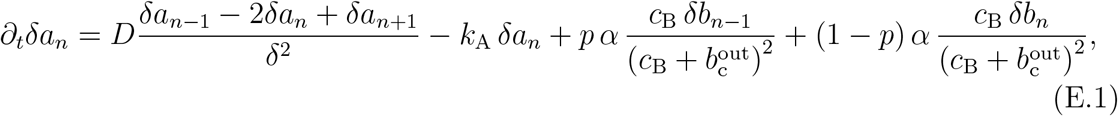

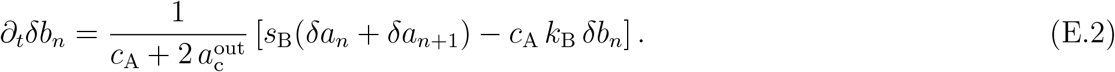

We use the ansatz

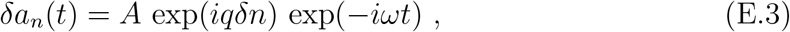

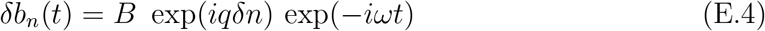

for the eigenmodes of the system, where *q* specifies an inverse wavelength, *δ* denotes the cell width, *ω* denotes a relaxation rate, *t* specifies the time, *n* the spatial index of the cell or extracellular space, and *A, B* ∈ 𝒞 denote the amplitudes of the eigenmodes that can be *q*-dependent. With this ansatz, the dynamics simplify to the eigenvalue problem

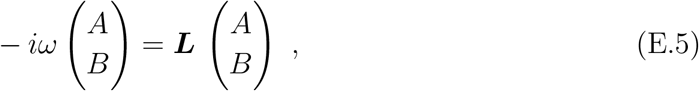

where

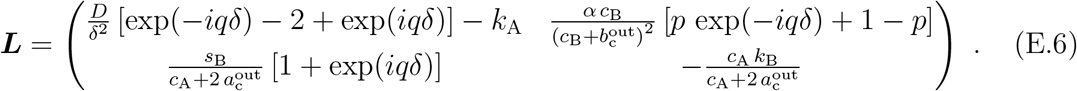

**Figure D1.**
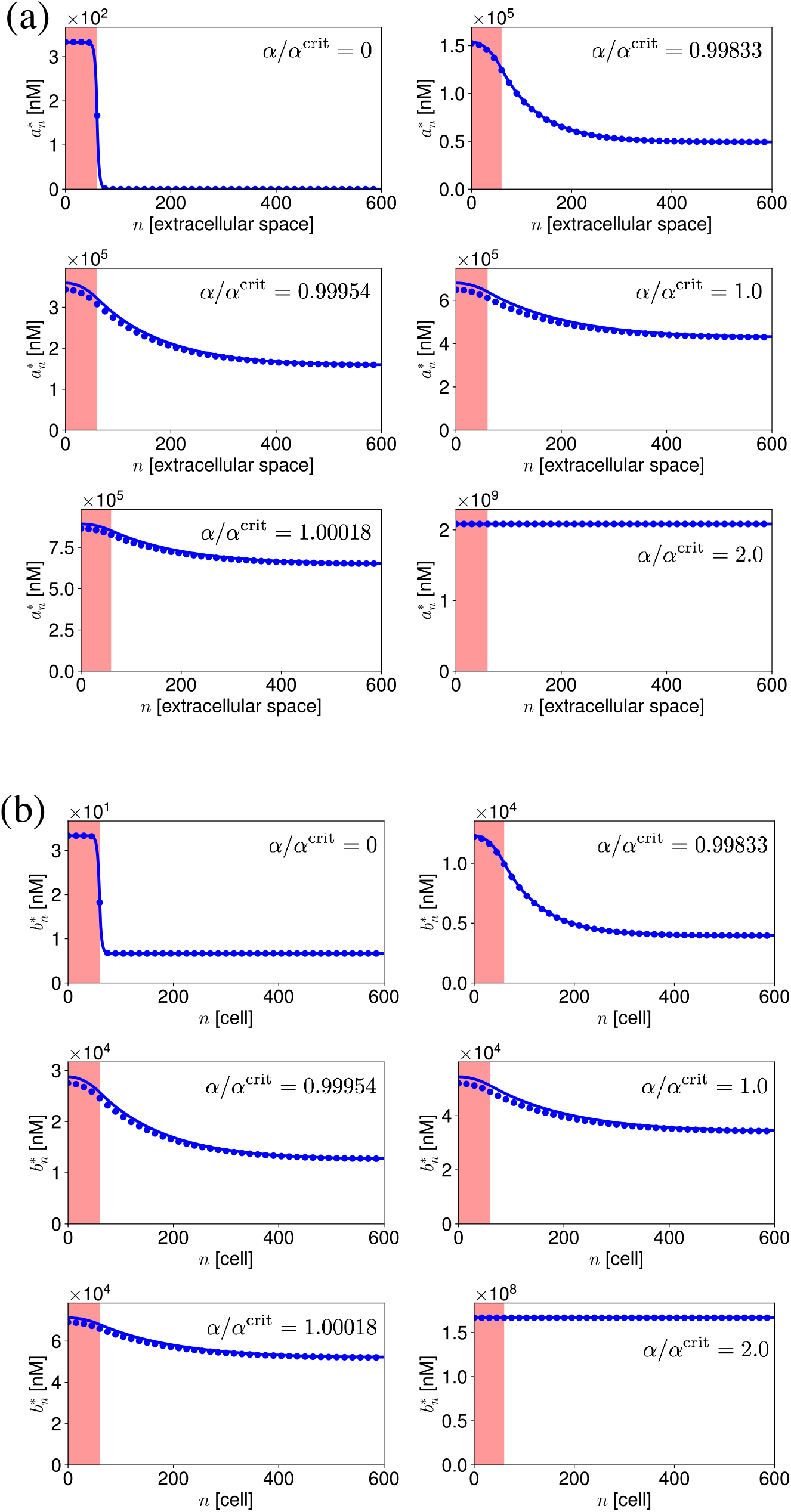
Steady-state concentration profiles of the morphogen (a) and the signaling activity (b) for indicated values of the feedback strength *α*. The signaling-independent source region is shaded in red. Circles denote the numerically obtained steady-state solution to the non-linear equations, lines indicate the analytical approximation to the steady state. Parameter set used for figures 2, 3.

**Figure D2.**
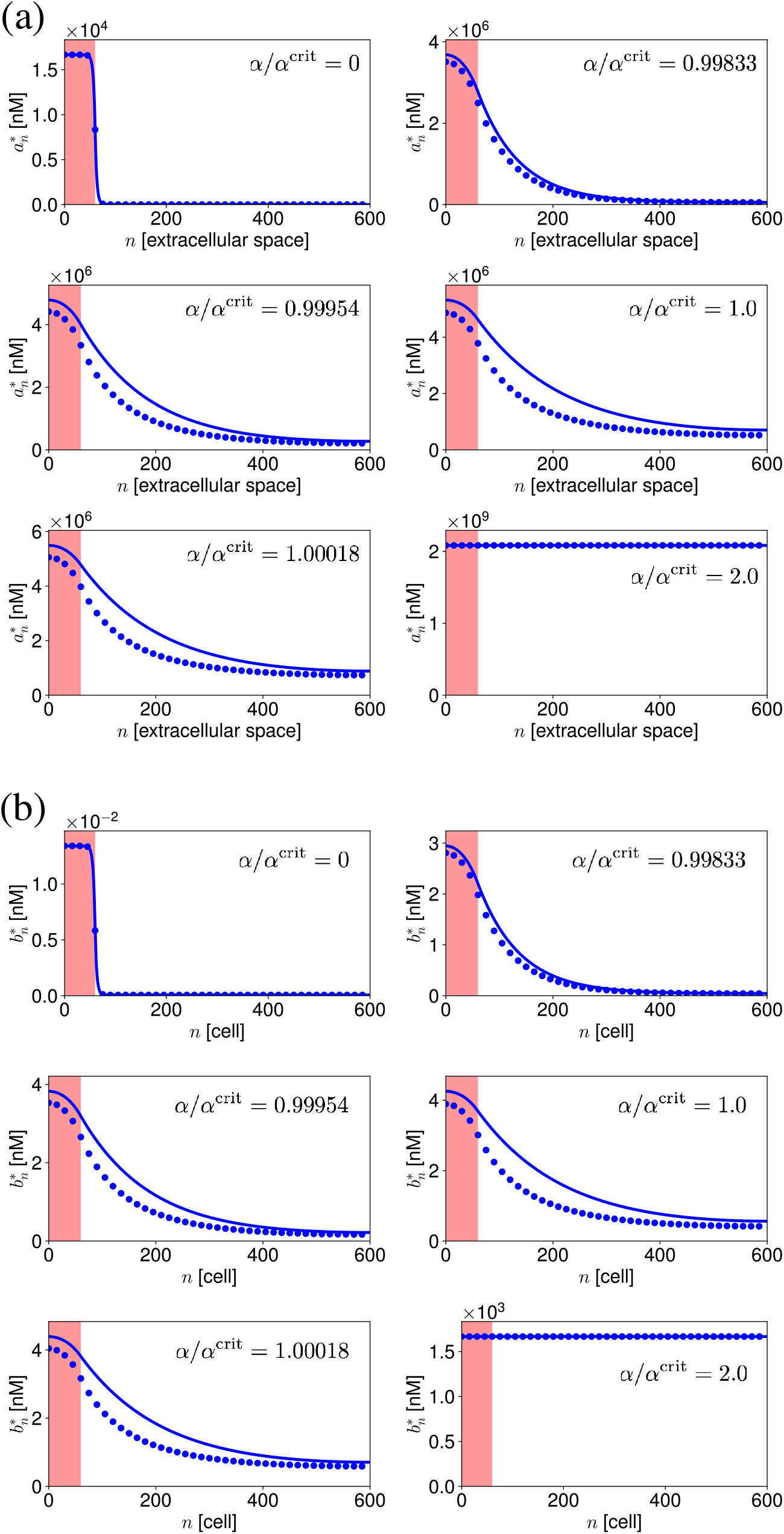
Steady-state concentration profiles of the morphogen (a) and the signaling activity (b) for indicated values of the feedback strength *α*. The signaling-independent source region is shaded in red. Circles denote the numerically obtained steady-state solution to the non-linear equations, lines indicate the analytical approximation to the steady state. Parameter set used for figures 4 -7, p = 0.50.

From the resulting characteristic polynomial, we obtain two relaxation rates *ω*_1_ and *ω*_2_ which depend on the wave number, i.e. the inverse wavelength *q*:

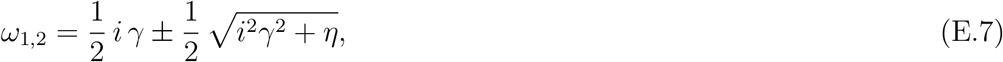

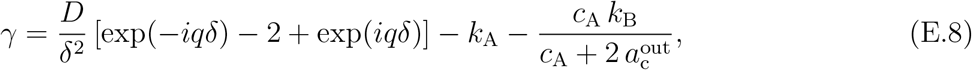

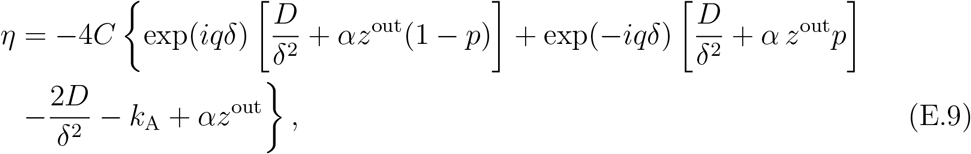

where *z*^out^ is defined by equation (12) and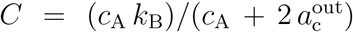. We aim to analyze the dynamics in the vicinity of the steady state. The steady state of the system is contained in these dynamics as the limit of *ω* = 0. We see that 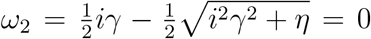 if *η* = 0. Thus, the steady-state and therefore the slow dynamics are contained in the relaxation rate *ω*_2_. We hence focus our further analysis on this relaxation rate. We identify the two roots of *ω*_2_ determined by *η* = 0. We start by expanding the exponential functions for small values of *iqδ*, i.e. long wavelengths, using

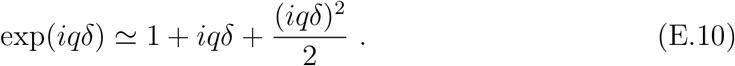

We obtain the approximations

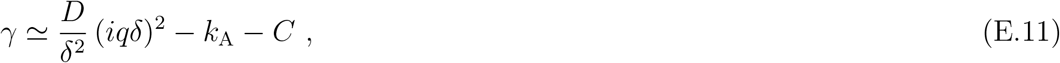

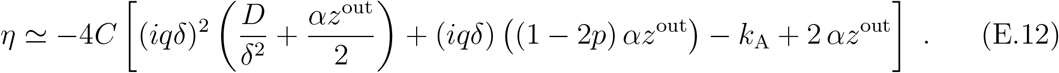

Based on these, we can then calculate the inverse steady-state decay length *iq*_1,2_*δ* defined by the steady-state condition *η* = 0. We obtain

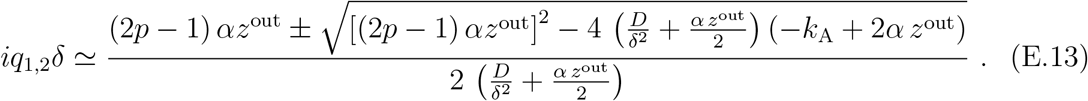

Note that *q*_1,2_ are equivalent to the steady-state decay lengths obtained in Appendix D, equations (D.17), (D.18):

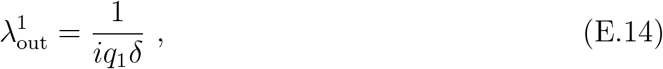

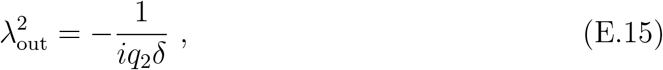

where the difference in sign is due to the definition of signs in equation (6). We can then expand *ω*_2_ around its roots, i.e. the inverse steady-state decay lengths *q*_1,2_. To this end, we define

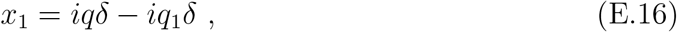

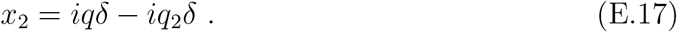

We can then express *η* as

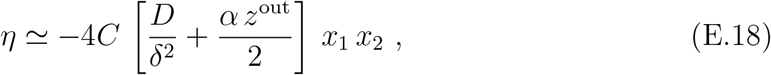

and *γ* as

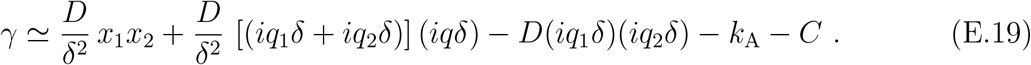

Note that the steady state now corresponds to *x*_1_ = 0 or *x*_2_ = 0. We aim to expand *w* for wavenumbers in the vicinity of the steady-state, i.e. small *x*_1_*x*_2_. The hydrodynamic limit can be further approximated considering *x*_2_ small such that *iqδ* ≃ *iq*_2_*δ*. With these simplifications, *γ* can be approximated as

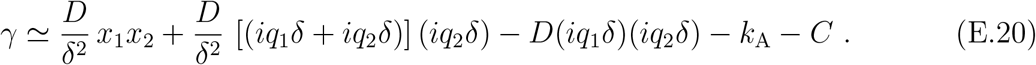

Using equations (E.16) - (E.20), we can then expand *ω*_2_ to lowest order in *x*_1_ *x*_2_ as

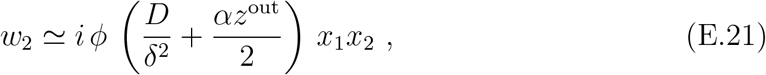

where

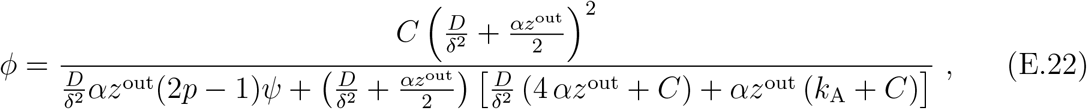

where

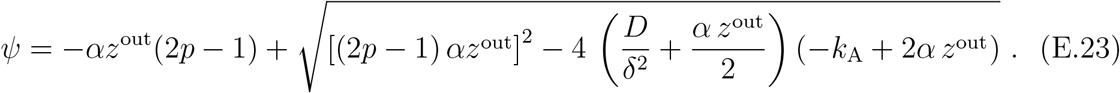

With

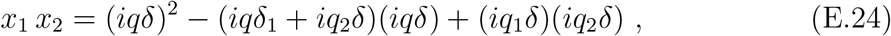

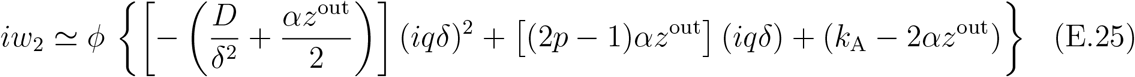

equation (E.25) can be compared to the dispersion-relation of a convection-diffusion-degradation equation (equation (8)) which reads

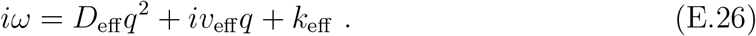

By comparing equation (E.25) and (E.26), we can identify the effective transport coefficients and loss rate as the coefficients of powers of *q*. They are given in equations (9) - (11).

### Appendix F. Decay length based on numerical steady-state solution

In order to compute the decay length based on the numerical steady-state solution, we assume that the profile outside of the source region follows an exponential decay added on to a constant plateau or baseline. To reveal its decay length, we numerically compute the second derivative of the numerical steady-state profile. We obtain a base-line corrected numerical steady-state profile by subtracting the base-line or plateau value (given by the value at the distal end of the numerical steady-state profile) from all points of the profile. Subsequently, we point-wise divide this base-line corrected profile by the second derivative of the profile. We take the square root of these values and use their median of in the non-source region as the decay length based on the numerical steady-state profile.

### Appendix G. Relaxation times

In order to obtain the relaxation modes of the signaling relay, we analyze the dynamics of the system close to the steady state (see equations (19), (20)). We approximate the dynamics of the system close to the steady state by linearizing the dynamic equations of the morphogen (equation (1)) and the intracellular signaling molecule concentration (equation (2)) around the steady-state solution of the system given by 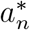 and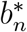. We obtain:

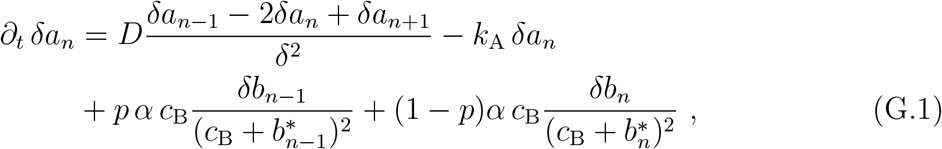

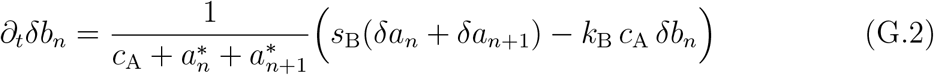

in the bulk of the system. The linearized dynamic equations at the boundaries can be obtained in the same way.

We can obtain the steady-state profiles 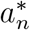 and 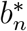 numerically, see Appendix C. Alternatively, we can use the approximation to the steady-state profiles discussed in section 3.1, equation (6), using equations (5) and (6) as 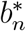 and 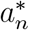, respectively. The approach for analyzing the dynamics stays the same, the only difference is which profiles are used to linearize the dynamics.

Either way, the resulting system of linear equations can be expressed in matrix form as:

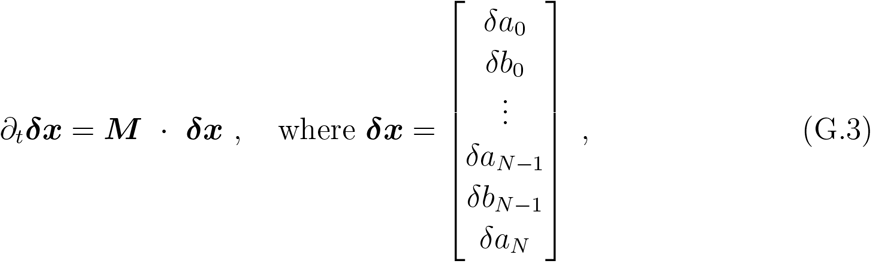

and ***M*** describes the dynamics of the system according to equations G.1 and G.2 and the respective linearized equations at the boundaries. The dynamic solution to this system is given by the sum of all eigenmodes:

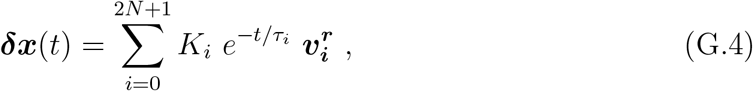

where 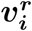 denotes a right eigenvector of ***M*** with eigenvalue 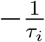, and the *τ*_*i*_’s denote the relaxation times of the system. We need to distinguish between the left and the right eigenvectors, since M is not symmetric and hence the right eigenvectors are not necessarily orthogonal to their transpose. Note that the system is comprised of both the cells and the extracellular spaces, gathered in ***δx***, and thus has one set of 2 *N* + 1 relaxation times and corresponding eigenvectors. We obtain the eigenvectors and eigenvalues by numerically diagonalizing ***M***. The coefficients *K*_*i*_ are determined by the initial conditions, i.e. the initial concentration of morphogen (***a***(0)) and the initial concentration of the intracellular signaling molecule (***b***(0)). Gathering these initial conditions, as well as the steady-state solution in respective vectors similar to what we did for *δx*:

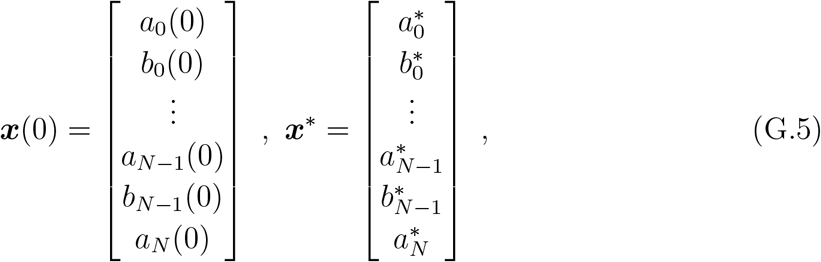

we obtain the *K*_*i*_ according to:

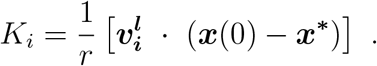

That is, we compute the *K*_*i*_ by projecting the initial condition onto the corresponding elements of the left eigenvector 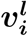 and normalizing to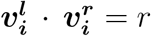. Note that we use the *left* eigenvectors to compute the *K*_*i*_ as we used the *right* ones to obtain the eigenmodes.

Based on this approximation to the dynamic solution of the system close to the steady state, we can define the characteristic time scale of the relay mechanism. For a linear system like the one presented in equation (G.3), the dynamics of the system are governed by the slowest relaxation time *τ*_max_ for long times. In particular, this slowest relaxation time is a good estimate of how long it takes the linear system to reach its steady state. We thus define the slowest relaxation time

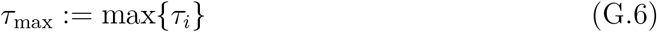

as the time scale of the relay mechanism.

### Appendix H. Parameters used for figures

Table H1 summarizes the parameters used in the figures.

**Table H1.** Parameters used for figures.

| Parameters | Figure 2 | Figure 3 | Figure 4 | Figure 5 | Figure 6 | Figure 7 |
| --- | --- | --- | --- | --- | --- | --- |
| $\alpha$ [(s V)<sup>-1</sup>] | $0.99833 * \alpha^{\text{crit}}$ | varying | varying | varying | varying | varying |
| $\alpha$ [(s V)<sup>-1</sup>] | $0.99833 * \alpha^{\text{crit}}$ | varying | varying | varying | varying | varying |
| $c_A$ [nM] | $1.66 \times 10^2$ | $1.66 \times 10^2$ | $1.66 \times 10^2$ | $1.66 \times 10^2$ | $1.66 \times 10^2$ | $1.66 \times 10^2$ |
| $c_B$ [nM] | $1.66 \times 10^8$ | $1.66 \times 10^8$ | $1.66 \times 10^3$ | $1.66 \times 10^3$ | $1.66 \times 10^3$ | $1.66 \times 10^3$ |
| $D$ [$\mu\text{m}^2 \text{ s}^{-1}$] | 1.0 | 1.0 | 1.0 | 1.0 | 1.0 | 1.0 |
| $\delta$ [$\mu\text{m}$] | 10.0 | 10.0 | 10.0 | 10.0 | 10.0 | 10.0 |
| $k_A$ [s<sup>-1</sup>] | $10^{-3}$ | $10^{-3}$ | $10^{-3}$ | $10^{-3}$ | $10^{-3}$ | $10^{-3}$ |
| $k_B$ [s<sup>-1</sup>] | $10^{-3}$ | $10^{-3}$ | 100 | 100 | 100 | 100 |
| $N$ | 600 | 600 | 600 | 600 | 600 | 600 |
| $s_A$ [molecules (s V)<sup>-1</sup>] | 20 | 20 | 1000 | 1000 | 1000 | 1000 |
| $s_B$ [molecules (s W)<sup>-1</sup>] | 4 | 4 | 4 | 4 | 4 | 4 |
| $p$ | 0.5 | 0.5 | see legend | see legend | see legend | see legend |
| $V$ [$\mu\text{m}^3$] | $10^2$ | $10^2$ | $10^2$ | $10^2$ | $10^2$ | $10^2$ |
| $W$ [$\mu\text{m}^3$] | $10^3$ | $10^3$ | $10^3$ | $10^3$ | $10^3$ | $10^3$ |
| $w$ | 60 | 60 | 60 | 60 | 60 | 60 |

